# The impact of short- and long-range perception on population movements

**DOI:** 10.1101/440420

**Authors:** S. T. Johnston, K. J. Painter

## Abstract

Navigation of cells and organisms is typically achieved by detecting and processing orienteering cues. Occasionally, a cue may be assessed over a much larger range than the individual’s body size, as in visual scanning for landmarks. In this paper we formulate models that account for orientation in response to short- or long-range cue evaluation. Starting from an underlying random walk movement model, where a generic cue is evaluated locally or nonlocally to determine a preferred direction, we state corresponding macroscopic partial differential equations to describe population movements. Under certain approximations, these models reduce to well-known local and nonlocal biological transport equations, including those of Keller-Segel type. We consider a case-study application: “hilltopping” in Lepidoptera and other insects, a phenomenon in which populations accumulate at summits to improve encounter/mating rates. Nonlocal responses are shown to efficiently filter out the natural noisiness (or roughness) of typical landscapes and allow the population to preferentially accumulate at a subset of hilltopping locations, in line with field studies. Moreover, according to the timescale of movement, optimal responses may occur for different perceptual ranges.

## 1. Introduction

Navigation and migration play numerous crucial roles, for example allowing circulating immune cells to seek and destroy infections, and populations to collect at breeding grounds. A key ingredient for successful navigation lies in the availability of external orienteering cues, ranging from chemicals to visual landmarks, interpreted via (for organisms) a variety of sensory organs (eyes, ears, noses, lateral lines *etc.*) or (for cells) highly specific transmembrane receptors. The means by which cues are detected, processed and integrated are, naturally, subject to considerable speculation [14].

Numerous models have been formulated to describe oriented movement, most frequently for chemical gradient responses (chemotaxis). Given that movement data is typically recorded at an individual level (e.g. cell/organism tracking), a natural approach is to model an individual’s movement as a velocity-jump random walk (VJRW) [32]. In a VJRW, an individual undergoes an alternating sequence of smooth “runs” with constant velocity and “turns” where a new velocity is chosen. We note that the latter provides a point for including orientation. In a groundbreaking paper, Patlak [37] derived a governing partial differential equation (PDE) for motions of this form, a model of advection-diffusion type in which an external bias drifts the population in a specific direction. This model foreshadowed phenomenological models such as the well-known Keller-Segel [22] model for chemotactic guidance, a popular choice for incorporating taxis-type orientation into models for biological movement [35].

Many models implicitly assume that guidance information is local to the individual’s position. For an adult fruitfly, capable of detecting and orienting according to a (spatial) chemical gradient computed across its antennae [6], the length scale of antenna separation is microscopic in comparison to its movement in the landscape at large. The scale of gradient detection is, effectively, local. Yet in other cases the sampling region for cue detection may be much larger, covering a significant portion of the individual’s environs. An obvious example lies in visual sensing, where individuals can assess information within eyesight. Here, the *perceptual range* - the “distance from which a particular landscape element can be perceived as such (or detected) by a given animal” [24] – is a key concept and has been estimated at distances ranging from metres to kilometres, according to species, habitat and conditions [54, 43, 30, 51, 11, 46, 12]. Auditory cues are also detectable at a distance: a few kilometres for elephants [26] and possibly hundreds of kilometres for baleen whales [38], the latter possible via the properties of sound propagation in water. Less obviously, fish and amphibians detect the electric fields generated by nearby organisms and objects, a process termed electrolocation [20]. Long-range assessments can also extend down to the single cell, with certain cells generating membrane protrusions (filopodia, cytonemes) that stretch multiple cell diameters to acquire inherently nonlocal information about their tissue landscape [50, 23].

In this paper we explore models, founded on a simplified VJRW, that incorporate local and nonlocal evaluation of the environment. Orientation enters by biasing turns into certain directions, according to an underlying *navigation field* that can be adapted to account for various modes of sampling the environment. As prototype models we consider three simple responses to a scalar cue: a local gradient-based orientation, a nonlocal gradient-based orientation and a nonlocal response in which orientation is according to the maximum cue value over the perceptual range. The corresponding macroscopic models for the VJRW are of drift-anisotropic diffusion type. Via approximating expansions, we show how the simple local gradient model reduces to a Keller-Segel equation while the nonlocal models reduce to forms related to commonly-employed integro-PDE equations, such as those used to describe nonlocal chemotaxis responses [17], cell interactions [36], swarming/flocking [28] and, pertinently, information gathering over some perceptual range [10]. While these simplified forms strictly apply only when the orientation is weak, higher order approximations can extend the range of application. Separate approximations can be applied for very strong orientation responses, generating a pure drift equation in the simplest case.

To demonstrate how nonlocal assessment contributes to population structuring, we consider a specific case-study application: hilltopping in butterflies, moths and other insects. In hilltopping, population members move up slope and localise at peaks and summits, a “lekking” behaviour that increases mating encounters [48, 47, 1]. Given that elevation is (primarily) sensed visually, we explore how perceptual range and assessment mode impacts on the macroscopic population movement. Gradually introducing noise into an idealised elevation profile, we show how nonlocal responses can allow a population to overcome roughness. Using terrain data for the Bodega Marine Reserve, a recent site of field studies [15], these findings are shown to extend to natural environments, with nonlocal sampling allowing the population to accumulate on a select subset of prominent peaks. We discuss the results both generally and within the specific context of hilltopping.

## 2. Model framework

We first outline the framework, referring to [32, 45, 18, 52] for further details. We let *t* denote time, **x** ∈ ℝ^*l*^ the spatial coordinate and **v** ∈ ℝ^*l*^ the velocity. While the following derivation is general, in this work we focus on two-dimensional navigation (*l* = 2). This type of navigation is relevant for movement across a landscape. Individuals move via a VJRW with instantaneous waiting time between reorientations, requiring two probability distributions: a runtime distribution over ℝ+ that dictates the reorientation rate and a turning distribution over velocity space *V* that describes the new velocity. The former is taken to be a standard Poisson process, that is, exponentially distributed runtimes with constant mean runtime *τ* (or turning rate *λ* = 1/*τ*). We note that other choices of runtime can be made, potentially giving rise to subdiffusive or superdiffusive behaviour [13, 49, 9]. For the turning distribution we assume: (i) individuals move with a fixed speed *s* (i.e. **v** = *s*𝕊^*l*−1^); (ii) the new heading does not depend on the previous heading. Assumption (i) means that the turning distribution can be defined in terms of a directional distribution on the unit sphere, *q*(**n**|*t*, **x**), specifying the probability of choosing direction **n**∈𝕊^*l*−1^ following a turn, where **n**is the directional heading on the unit sphere 𝕊^*l−*1^.

For the above VJRW, given stochastically independent walkers, we can obtain an evolution equation for the macroscopic population density at position **x** and time *t*, *υ*(**x**, *t*). The process is to first write down the analogous transport (or kinetic) equation and, via scaling, obtain a macroscopic equation in an appropriate limit. We refer to [18] (and references therein) for details and simply state that applying a moment closure method [^1^ Our moment closure relies on two frequently used assumptions: (i) the population’s variance is computable from the equilibrium distribution, and (ii) a fast flux relaxation. The latter effectively states that, at the space/time scales of interest, the particle responds instantaneously to the information obtained at its current position. In later simulations we compare directly the distributions from repeated simulations of the stochastic model and the macroscopic model, confirming their validity for the present problem.] generates a macroscopic model in the form of the Drift-Anisotropic Diffusion (DAD) equation

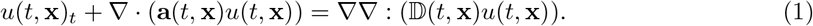

In the above, 𝔻(*t*, **x**) is a *l*×*l* diffusion tensor matrix and **a**(*t*, **x**) is the *l*-dimensional advection velocity. The colon (:) denotes the contraction of two tensors, resulting in a summation across all second order derivatives

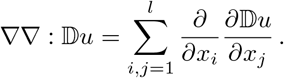

The macroscopic quantities **a** and 𝔻 are determined from the statistical properties of the individual-level model: mean speed, mean runtime and the turning distribution. Specifically,

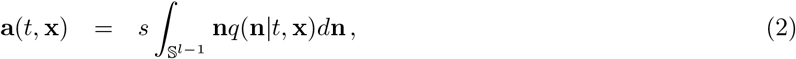

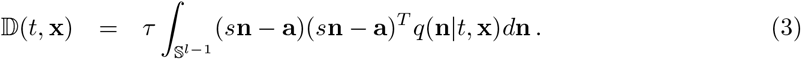

Thus, the drift is proportional to the expectation of *q* and the anisotropic diffusion is proportional to its variance-covariance matrix.

Environmental guidance is encoded in *q*, via biasing turns into specific directions. For the two-dimensional case considered here, the von Mises distribution [25] is a natural choice:

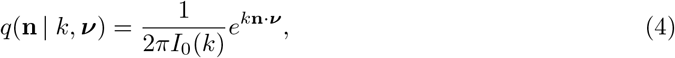

where *I*_*j*_(*k*) denotes the modified Bessel function of first kind of order *j*. Equation (4) plays an analogous role to the normal distribution of linear statistics, and generates a predominance of turns into *preferred direction* ***ν***∈𝕊^1^ with certainty increasing with *concentration parameter k*. We note that ***ν***(**x**, *t*) and *k*(**x**, *t*) are functions of space and time to reflect variation in the direction and strength of the orienteering cue, but we omit the dependences for clarity of presentation. As *k*→0, *q* tends to a uniform distribution, while as *k* → ∞, *q* approaches a singular distribution in which the preferred direction is always selected.

Expectations and variance-covariance matrices of (4) can be explicitly calculated [19], yielding the following drift-velocity and diffusion tensor

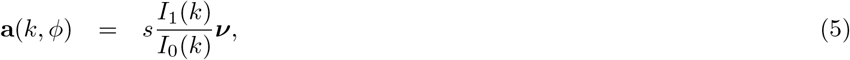

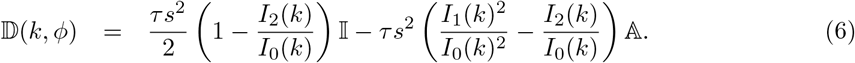

Intuitively, the drift is in the dominant direction. The anisotropic diffusion tensor has been decomposed into isotropic (proportional to the identity matrix 𝕀) and anisotropic (proportional to the singular matrix 𝔸 = ***νν***^*T*^) components. The relative strength of diffusion to advection depends on *k*, via the modified Bessel functions. We note that

- 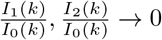 as *k* → 0 and we converge to isotropic diffusion;
- 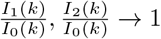 as *k* → ∞ and we converge to a pure-drift equation;
- 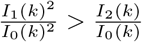 for all *k*, and hence the dominating axis of diffusion lies orthogonal to that of drift.

Modified Bessel functions arise in numerous applications and a correspondingly large literature has emerged [31]. In particular, approximating expansions are available for small and large arguments, as presented in Appendix A.

## 3. Orientation: local and nonlocal sampling

Orienteering information is encoded into a *navigation vector field*, denoted **w**(**x**, *t*) ∈ ℝ^*l*^. The directions and vector lengths of **w** form the inputs for the von Mises distribution: 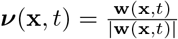 and *k*(**x**, *t*) = |**w**(**x***, t*)|. The navigation field in turn depends on a navigating factor, represented as a single scalar intensity function *E*(**x**, *t*), such as chemical concentration, elevation, temperature or magnetic intensity. We define **w**to have one of two generic forms:

- *Local sampling* – the navigation field is computed according to information obtained strictly at an individual’s current position;
- *Nonlocal sampling* – the navigation field is computed over some spatially-extended sampling region, centred on the current position.

**Figure 1:**
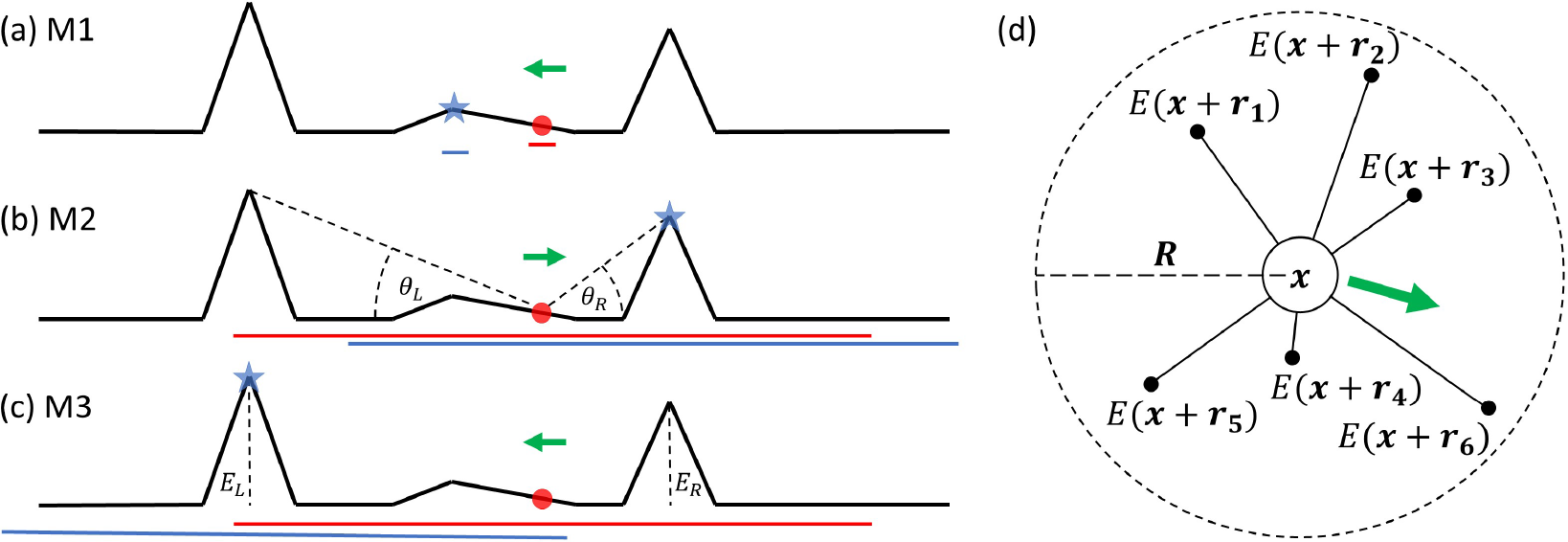
(a-c) Illustration of (M1)-(M3) in the context of hilltopping behaviour. Red circles and blue stars indicate starting and expected final locations of the individual; green arrows represent the initial direction of the bias; red and blue lines indicate the sampling region at the start and at the end of movement. (a) Local slope response; (b) Individual responds to the elevation angle, with *θ*_*R*_ > *θ*_*L*_ implying a bias to the right; (c) Individual responds to the absolute elevation, with *E*_*L*_ > *E*_*R*_ implying a bias to the left. (d) Illustration of nonlocal sampling. An individual at **x** samples the environmental cue *E* at points within some compact region (here taken to be a circle centred on **x** and characterised by the perceptual range, *R*), averaging the response into a preferred directional bias (green arrow).

For both sampling mechanisms we consider a general form before tailoring into the following three model prototypes:

(M1) a local and linear gradient model;
(M2) a nonlocal and nonlinear gradient model;
(M3) a nonlocal and nonlinear maximum model.

The above have widespread applicability, but are particularly intuitive in the context of hilltopping. In (M1), responses are to the *local gradient*, as highlighted in Figure 1(a). An individual, initially at the red circle, moves up the slope until coming to rest at the nearest local peak (blue star). In (M2), presented in Figure 1(b), evaluations are made up to a given perceptual range (red/blue lines). The bias is weighted in the direction of the *maximum elevation angle* from the current position and, while all three peaks are observed from its initial position, the individual typically moves towards the closer of the two higher peaks: despite its lower overall height, it has the larger elevation angle from the initial location. In (M3), the individual estimates the *true height* of peaks and biases according to the highest peak within perceptual range, as highlighted by the schematic in Figure 1(c).

### 3.1. Local sampling

Local models implicitly assume pointwise sampling (at the scale of the overall environment). To generate a directional response we assume the local gradient of *E* can be determined. This results in a (local) tropotaxis response in which **w** points in the direction of ∇*E* and the magnitude depends on *E* and |∇*E*| (and other factors, if necessary). Thus,

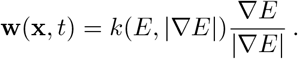

where *k*(*E*, ∇*E*) describes the strength and form of response: positive (negative) values of *k* imply attraction (repulsion). We note that *k*(*E*, ∇*E*) must satisfy *k* = 0 for |∇*E*| = 0. Given data, *k* can be estimated through fitting, for example in leukocyte chemotaxis where chemokine responses have been calculated for different concentrations and gradients [21].

To generate the prototype model (M1) we invoke the simple choice of linear gradient dependence: *k* = *κ*_1_|∇*E*| with associated response coefficient *κ*_1_. The navigation field is therefore directly proportional to ∇*E*, yielding

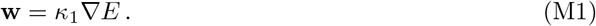

We remark that similar models have been proposed in a cell biology context, where *E* arises from local crowding between cells [3]. Movement in these models can either be repulsive and hence proportional to −∇*E*, or attractive and hence proportional to ∇*E*[3]. We note that such models consider navigation based from interactions between individuals, whereas we focus on processes where individuals do not interact with each other. Instead, navigation information is obtained from an underlying environmental cue.

### 3.2. Nonlocal sampling

We consider the schematic in Figure 1(d) and suppose an individual at **x** computes a movement response by directly sampling the environment at positions **x** + **r**_*i*_ for *i* = 1…*L*, where *L* is total number of perceived locations. Sampling could occur through (for an organism) visual focus on a distant point or (for a cell) extending a filopodium. We define a probability distribution, Ω(**r**|**x**, *t*), that denotes the probability that position **x** + **r** is sampled from position **x** at the time of reorientation *t*. Logically, Ω has compact support to reflect a maximum perceptual range, but analytically convenient forms with decaying tails (such as Gaussians) may also be reasonable. We suppose the strength of attraction towards (or repulsion from) **x** + **r** is encoded into a response function *g*(*E*(**x**+**r**, *t*), *E*(**x**, *t*), |**r**|), potentially depending on the cue at both the current and sampled points, as well as the sampling distance. The attraction vector **p**_**r**_(**x**, *t*) in direction **r**/**r** due to information at **x** + **r** is hence

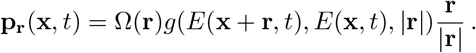

We suppose **w** is formed from the net attraction, obtained by sampling over different locations and calculating the average. Thus, we take

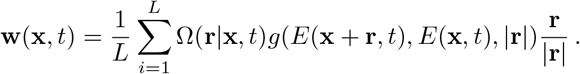

If a large number of points are sampled we can approximate **w** via the integral form

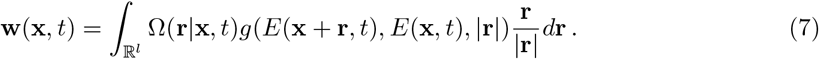

This generic formulation is easily adapted and we next consider the simple prototype models. Note that here, for simplicity, we exclusively concentrate on uniform sampling over a circular region of radius *R*, where we define *R* to be the *perceptual range*

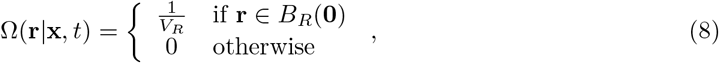

where *B*_*R*_(**0**) is the ball of radius *R*, centred on the origin, and *V*_*R*_ is its volume (i.e. *V*_*R*_ = *πR*^2^ for *l* = 2).

#### 3.2.1. Nonlocal gradient sampling

Suppose attraction is according to the *nonlocal gradient* of *E*

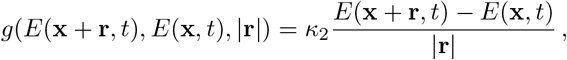

with sensitivity coefficient *κ*_2_. Substituting both the above and (8) into (7), we obtain

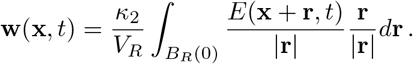

Notably, a Taylor series expansion about **x** applied to the RHS yields

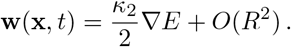

Hence, as *R* → 0 the nonlocal gradient model converges to the local gradient model with *κ*_2_ = 2*κ*_1_.

It is straightforward to incorporate nonlinear processing of the cue, and in particular we consider

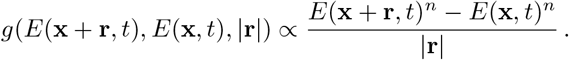

The parameter *n* gives additional control, increasing the weighting of attraction bias as *n* is increased. This gives the following navigation field (M2)

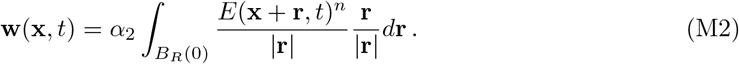

We note that we have absorbed various constants into 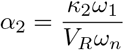 where

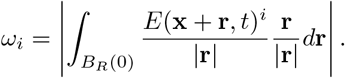

The scaling normalises the navigation field, so that the magnitude remains essentially the same as *n* is altered. Note that Taylor expansion again yields (M1) in a small *R* approximation.

#### 3.2.2 Maximum value evaluation

Our second prototype nonlocal model takes *g* to depend on the absolute size of the cue at *E*(**x**+**r**,*t*)

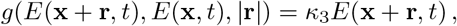

where *κ*_3_ is the sensitivity coefficient. Substituting into (7) along with (8) now yields

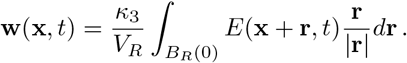

The distinction with the gradient-based evaluation is exemplified through a Taylor expansion, where we find

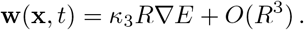

Hence, as *R* → 0 the navigation vector field shrinks to zero length and orienteering information is lost. As before, we can extend to a nonlinear form to obtain (M3)

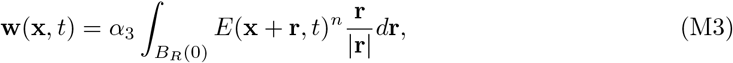

where 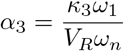 and

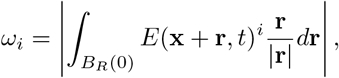

provides the normalisation as *n* is altered.

## 4. Approximations

A choice must be made about whether to perform direct stochastic simulations of the VJRW model or solve the full DAD model (1) with their Bessel function dependent coefficients (5)-(6). Both approaches have associated costs. Simulations of the stochastic VJRW are expensive, scaling with population size; evaluations of the modified Bessel functions and implementing anisotropic diffusion terms are also costly. In an analytic context, anisotropic diffusion terms may be more difficult to deal with than isotropic counterparts. Consequently, it is worthwhile investigating simplifying reductions.

### 4.1. Local sampling approximations

We consider model (M1). Expansions of the modified Bessel functions (see Appendix A) allow a series of approximations to (1) to be obtained, relevant for small or large values of *k* (weak or strong navigating cues, respectively). For weak cues the lowest order approximation (*O*(*k*^1^)) yields the well known Keller-Segel equation for (chemo)taxis

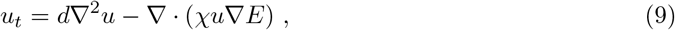

with constant diffusion (*d* = *τs*^2^/2) and chemotactic (*χ* = *sκ*_1_/2) coefficients [^2^ We remark the general choice *k* would instead yield a general form Keller-Segel equation in the first order approximation: 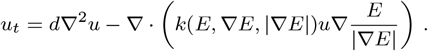]

The above is strictly valid only for weak cues, but a series of higher order “corrections” extends the range of *k* values for which the approximation accurately describes the full DAD model. We tabulate up to *O*(*k*^4^) in Appendix B. For example, the *O*(*k*^2^) approximation in two dimensions introduces a correction for each of the diffusion and chemotaxis terms, such that

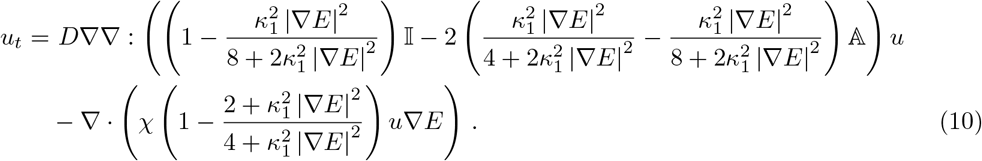

Notably, this correction introduces the anisotropic diffusion component. We remark that diffusion coefficients remain positive for all *κ*_1_, so the system is well-behaved.

As *k* becomes larger the approximations become less useful: more terms are required to achieve reasonable accuracy, and it would be natural to instead revert to the full DAD model (1). However, very large *k* can alternatively be dealt with through large argument approximations (Appendix B). For example, for sufficiently large *k* (corresponding to strong navigation fields) we can approximate the full DAD model (1) via the pure drift-equation

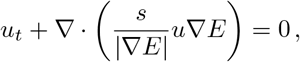

which simply generates direct movement with speed *s* in the direction of the steepest gradient. Corrections to this again yield a sequence of chemotaxis/anisotropic diffusion style models, tabulated in Appendix B. For example, the *O*(*k*−1) approximation in 2D generates

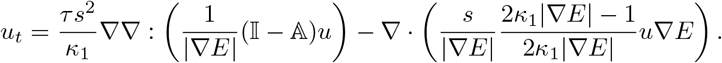

The anisotropic diffusion in the above corresponds to a degenerate/singular form in which diffusion is one-dimensional and along the axis orthogonal to the preferred direction.

### 4.2. Nonlocal sampling approximations

The process is identical for nonlocal sampling scenarios, and we therefore only state the first order weak cue form for the general navigation field (7), and for both (M2)-(M3). In the general case, the first order approximation under a weak cue yields a nonlocal integro-PDE equation

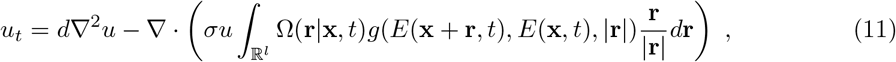

for *d* = *τs*^2^/2 and *σ* = *s*/2. The above is closely related to various nonlocal PDE models. For example, if *E* is directly based on the population’s own density distribution the above is similar to those employed to describe swarming [28] and cell interactions [36]. Moreover, the above is closely related to the model proposed in [10] for perceived gradient following.

If we consider the two prototype models, the above becomes

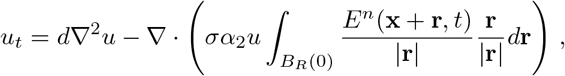

for (M2), and

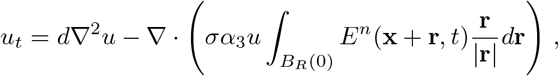

for (M3).

**Figure 2:**
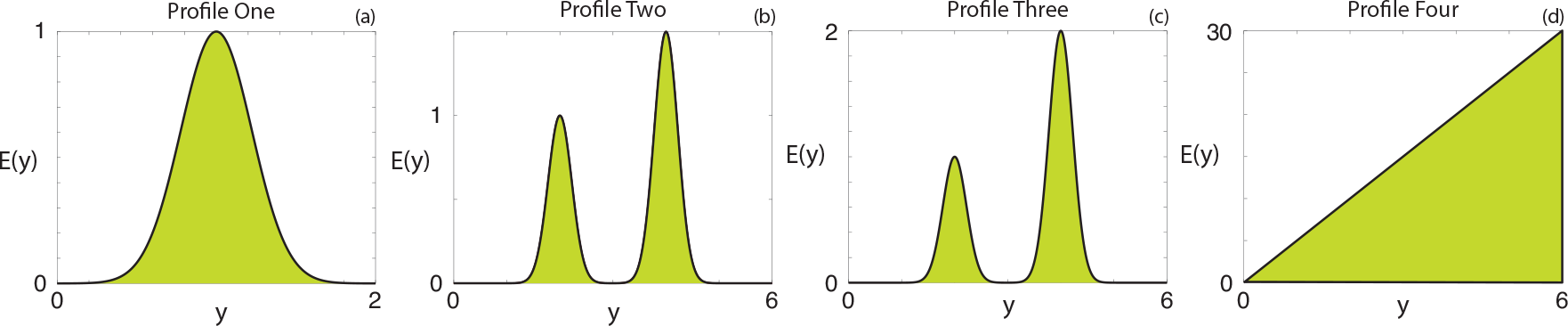
Four different idealised elevation profiles.

### 4.3. Numerical validation

We restrict our validation to local gradient sampling under an environmental cue that does not vary with time. The population is initially distributed uniformly across the rectangular region 0 ≥ *x* ≥ 1, 0 ≥ *y* ≥ 2. We generate a quasi-one-dimensional scenario by assuming the cue varies only with *y*, specifically

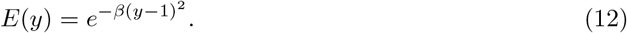

The corresponding cue profile is presented in Figure 2(a). The dominant direction for individuals with locations *y* < 1 (*y* >1) will be *π*/2 (−*π*/2) and subsequently individuals are expected accumulate along the ridgeline *y* = 1. Comparisons are presented for (i) the average behaviour of multiple realisations of the velocity-jump process, (ii) numerical solution of the full DAD model (1) under (5)-(6), and (iii) the *O*(*k*) to *O*(*k*^4^) approximations. These approximations rely on an assumption that the strength of the navigating cue, *k*, is small. Numerical methods are described in the Supplementary Materials, Section S3. Briefly, an implicit timestep method and an operator splitting method are used to numerically solve the full DAD model and the corresponding *O*(*k*) to *O*(*k*^4)^ approximations. The trajectories of individuals in the velocity-jump process are simulated directly as described in [34], and additionally detailed in the Supplementary Material, Section S3.

The Bessel function approximations are independent of *s* and we therefore expect that the difference between approximations and the full DAD model (1) is constant as *s* changes. Thus, differences between solutions under increasing *s* should stem only from the assumptions made when deriving the macroscopic model from the velocity-jump process. The difference between the approximations and either the full DAD model solution or the average behaviour of the velocity-jump process decreases with the order of the approximation. Further, both the full DAD model solution and approximations prove to be less accurate against the average velocity-jump behaviour as *s* increases (Supplementary Material, Section S1), consistent with previous investigations [34]. Generally, whether (1) provides a reasonable description of the underlying random walk depends on whether the spatial and temporal scales are appropriately macroscopic, for example see [5, 18] for further discussion. Specifically, average run lengths need to be short, compared to the overall domain dimensions, and individuals need to make numerous turns over the study time.

We next explore the match for fixed *s* = 0.001 and varying response coefficient *κ*_1_. Varying *κ*_1_ directly influences the concentration parameter *k*, so increasing *κ*_1_ should impact on the matching ability of the approximations. Note that for a quasi-one-dimensional case the diffusion tensor reduces to a function depending on *y* and *k*, denoted *d*(*y*), while the drift reduces to a “chemotaxis”-type function, denoted *a*(*y*). As expected, truncating at higher order terms yields a better fit for these functions, see Figures 3(a,d,g) for *a*(*y*) and Figures 3(b,e,h) for *d*(*y*). For sufficiently small *κ*_1_ (e.g. *κ*_1_ = 0.5) even the lowest order approximations prove highly acceptable, while larger values (e.g. *κ*_1_ = 2) demand a higher order for a good fit.

Turning to full numerical solutions, we compare *O*(*k*) to *O*(*k*^4^) approximations with those of the full DAD model and the average velocity-jump behaviour. Note that for the parameter values under consideration, the full DAD model closely describes the average velocity-jump behaviour. For *κ*_1_ = 0.5, Figure 3(c), all approximations are generally successful, although *O*(*k*) and *O*(*k*^2^) approximations show some inaccuracy in regions where the diffusivity function approximation is at its worst. Increasing *κ*_1_ generates a poorer fit: under this cue distribution and *κ*_1_ = 2, the maximum value of *k* ≈ 5.43 is beyond the expected range of *k* values for reasonably valid approximations. Nevertheless, the *O*(*k*^4^) still proves reasonably accurate.

Overall, numerical comparisons validate the approximations, but only when *k* remains within relevant regions. Thus, if it is known that a particular cue provides only a weak bias (say, a cell migrating in a fairly stable and shallow gradient) it is reasonable to use the simplified PDE, such as the local or nonlocal Keller-Segel equation. If, rather, cue strength cue varies widely, one should resort to employing the full DAD equation (1), assuming the spatial/temporal scales are appropriately macroscopic.

## 5. Short and long-range sampling: hilltopping case study

The accumulation of butterflies, moths and other insects at hilltops has been widely documented [48, 47, 8, 1, 43, 15]. Many insect populations fluctuate over different locations and seasons, and strategies that offer robustness against extinction events are necessary. Hilltopping is perceived as one such mechanism, due to an improved rate of mating encounters. The relatively local scale over which movements occur, the precision to which terrain can be documented and the (relative) ease of tracking/counting population members combine to form an ideal system for determining how environments structure a population. Given visual assessment of topography, it is natural to query how perceptual range impacts on hilltopping. Natural terrains are typically “rough”, so responses based strictly on the local gradient could easily trap a large percentage of the population, scattered through small accumulations on many local peaks. Instead, studies of a hilltopping butterfly (*Melitaea trivia*) indicate that their orientation may be according to both the local slope and higher peaks within a longer range (~50 metres) [43]. Moreover, a recent capture-mark-recapture study of the tiger moth (*Arctia virginalis*) revealed that populations accumulated only on a subset of summits, with the largest numbers found at the highest elevations [15].

Motivated by these findings, we use this section to explore how short- and long-range topographical sensing alters hilltopping outcomes. We consider our three prototype models for moving up slopes, effectively describing populations that respond to: (M1) the strictly local gradient; (M2) the nonlocal gradient, that is, according to the elevation angle of terrain within perceptual range; (M3) the true height of land within perceptual range. We first generate idealised terrains in order to test principles before considering an application using elevation data for the Bodega Marine Reserve, as used in a recent hilltopping survey conducted by Grof-Tisza *et al.* [15].

**Figure 3:**
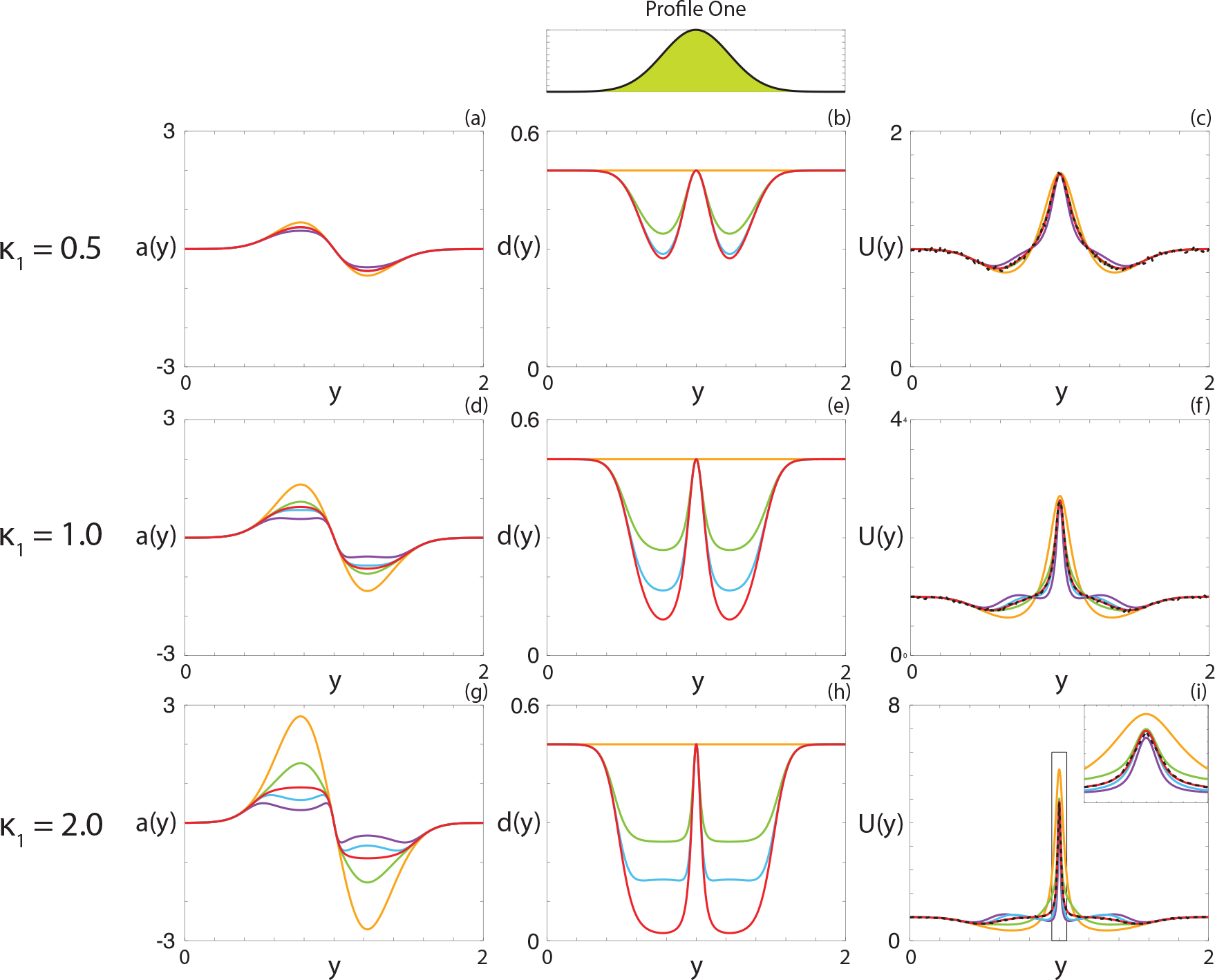
(a),(d),(g) Comparison between the drift-velocity function (red), and the corresponding *O*(*k*) to *O*(*k*^4^) approximations (orange, purple, green, blue, respectively) for (a) *κ*_1_ = 0.5, (d) *κ*_1_ = 1, (g) *κ*_1_ = 2. (b),(e),(h) Comparison between the diffusivity function (red), and the corresponding *O*(*k*) to *O*(*k*^4^) approximations (orange, purple, green, blue, respectively) for (b) *κ*_1_ = 0.5, (e) *κ*_1_ = 1, (h) *κ*_1_ = 2. (c),(f),(i) Comparison of the average velocity-jump behaviour (black, dashed), and the numerical solutions to the full DAD model (red) and the *O*(*k*) to *O*(*k*^4^) approximations (orange, purple, green, blue, respectively) for (c) *κ*_1_ = 0.5, (f) *κ*_1_ = 1, (i) *κ*_1_ = 2. Random walk parameters are set at *s* = 0.001 and *τ* = 1; elevation is set by (12) with *β* = 10; simulations are based on averaging over 10^6^ realisations of the VJRW and a spatial discretisation of Δ*y* = 0.001 for the numerical approximation of the continuous models. Simulations are computed until *t*_end_ = 100.

We note that several previous simulation studies have explored hilltopping. For example, in [41, 39] an agent-based modelling approach was used to simulate butterfly movement paths, based on field study data [43]. Topography was shown to channel movement paths along “virtual corridors”. Further agent-based studies in [42] highlighted the importance of an element of randomness in the movement paths to avoid the above described trapping. In [34] the multiscale framework used here was employed to demonstrated how accumulations formed by hilltopping could optimise mating for low-density populations, but may be disadvantageous for more abundant populations. The current study here extends that work, specifically incorporating long-range perception to explore its impact on population distribution.

### 5.1. Idealised study

We first explore how the prototype orienteering mechanisms impact hilltopping in a series of idealised studies. We assume a quasi one-dimensional terrain and, unless stated otherwise, initially suppose the population to be uniformly randomly distributed across the terrain. We first examine the suitability of models (M1)–(M3) to describe hilltopping, considering movement on a terrain containing a single peak centred at *y* = 1, as presented in Figure 2(a). For succinctness we simply note that all model prototypes allow the population to relocate to the peak by *t* = 100; results and further discussion are provided in the Supplementary Material, Section S2.1. Thus, all proposed mechanisms are plausible and we shift to a more sophisticated analysis of the mechanisms.

#### 5.1.1. Perceptual range and nonlocal response order

Here we investigate the influence of the perceptual range, *R*, and nonlocal response order, *n*. We consider a landscape containing two peaks of different heights, as highlighted in Figure 2(b). For model (M3) we observe a significant change between *R*= 1.5 and *R* = 2.0. For smaller radii the population is split into separated sub-populations, one on each hilltop with the majority of the individuals on the higher of the two peaks. In contrast, for *R* = 2.0 the individuals merge into a single contiguous population at the higher peak. Comparing the dominant direction for the two radii, Figure 4(c) and (f), a transition is observed such that under smaller radii each peak has its own “zone of attraction”. For larger radii, however, the zone of attraction for the larger peak expands to cover the entire domain. Note that for this example, *n* does not qualitatively change the results.

Under model (M2) with a high nonlocal order (*n* = 5), two distinct sub-populations are maintained even for large *R*. That is, even when the higher peak is fully within perceptual range of members on the lower peak, the two sub-populations remain separate. Here, higher weighting is applied according to the largest elevation angles. Consequently, the lower peak can always be perceived as more attractive for individuals located sufficiently close. As the nonlinear order is decreased, however, the sub-population that initially formed on the lower peak begins to migrate towards the higher peak. This is reflected in the comparison of the dominant direction, Figure 4(l), where we observe a collapse from two zones of attraction to one. As the nonlocal gradient model effectively places a weight on higher environmental cue values close to an individual, it is difficult for an individual to move away from the peak if that weighting is further compounded by a high nonlinear order.

To examine whether migration is unidirectional (from the lower to higher peak), we consider the migration proportion for two initially separated populations: one uniformly distributed around the lower peak and the second about the higher peak (see Supplementary Material, Section S2.1). Notably, when peak to peak migration occurs it is only for those initially located on the lower peak, which translocate to the higher one. We therefore investigate how the combination of perceptual range, ratio of peak heights and nonlocal order influences the proportion of the lower peak population that have migrated by a fixed time, set at *t* = 5000. The influence of peak height ratio is determined via two elevation profiles, Figures 2(b)-(c), with peaks located at the same position but with distinct heights to generate ratios of 1.5 and 2.0 respectively. Figure 5(a) summarises the results for (M3). For all scenarios, there is negligible migration for *R* = 1 and full migration by *R* = 2. In between, there is a steep transition with its location changing according to nonlocal order and peak height ratio: increases in either will lower the perceptual range required to shift the population. Effectively, an increase in either of these properties increase the attractiveness of the higher peak and the population migrates accordingly. Figure 5(b) summarises the results for (M2). Notably, there is now a significant difference between *n* = 1 and *n* = 5 for both profiles. As discussed previously, the extra weighting toward higher environmental values close to an individual, introduced by the 1/**r** factor in the nonlocal gradient term, influences the dominant direction (Figure 4(o)). Substantially altering the disparity between peak heights, though, can overcome this additional weighting.

**Figure 4:**
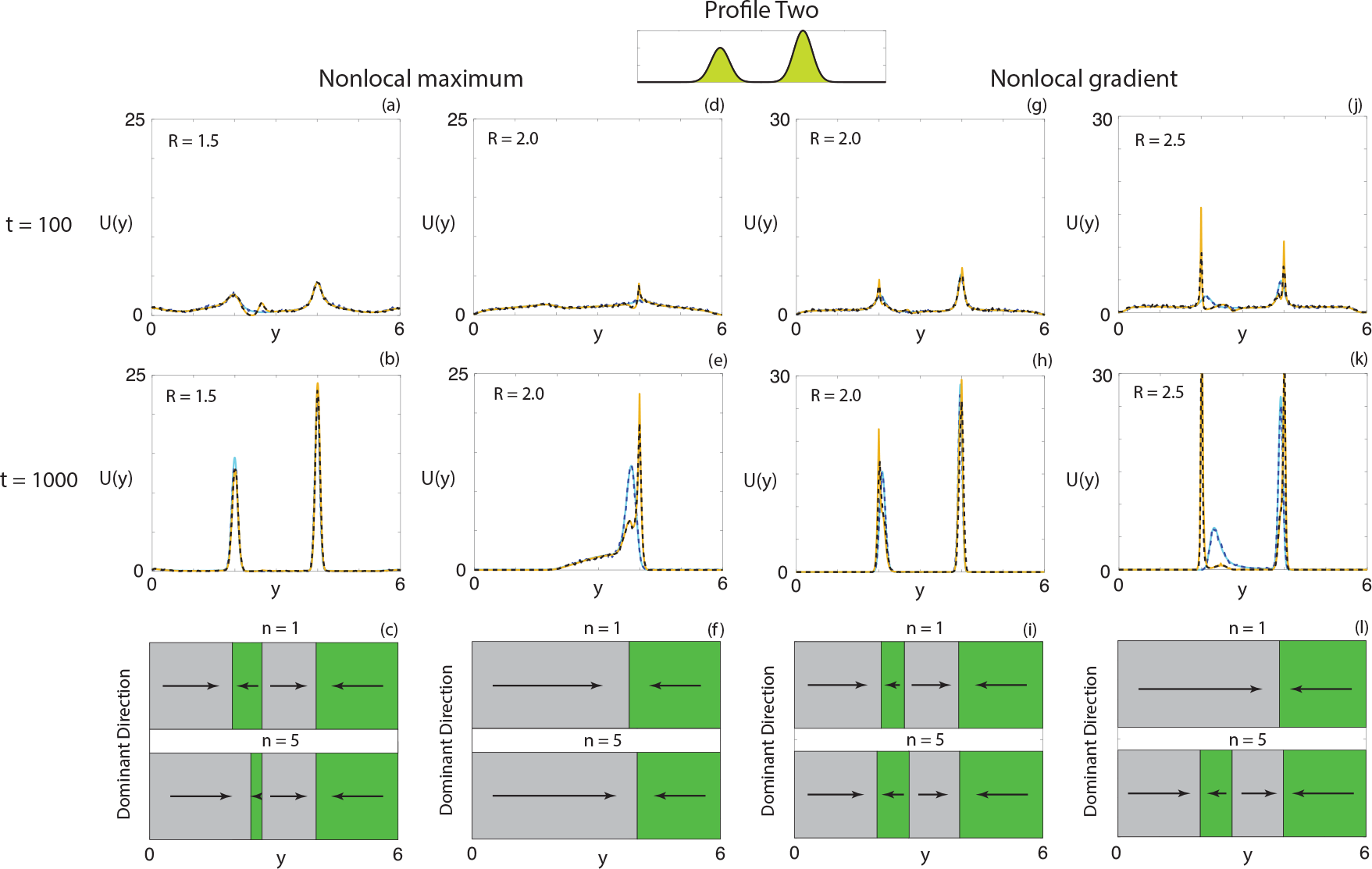
Comparison of different *R* and *n* for the full DAD model (solid) and the average velocity-jump process (dashed), respectively, using (a)-(f) (M3) and (g)-(l) (M2). Cyan/blue corresponds to *n* = 1, orange/black corresponds to *n* = 5. Initially the population is uniformly distributed and the elevation profile corresponds to Profile Two in Figure 2. Parameters used are as follows: (a)-(f) *κ*_3_ = 3, (g)-(l) *κ*_2_ = 0.3; random walk parameters are set at *τ* = 1, *s* = 0.01; numerical simulations based on averaging over 2 105 realisations of the VJRW and using a spatial discretisation of ∆*y* = 0.001 for the numerical approximation of the continuous models.

**Figure 5:**
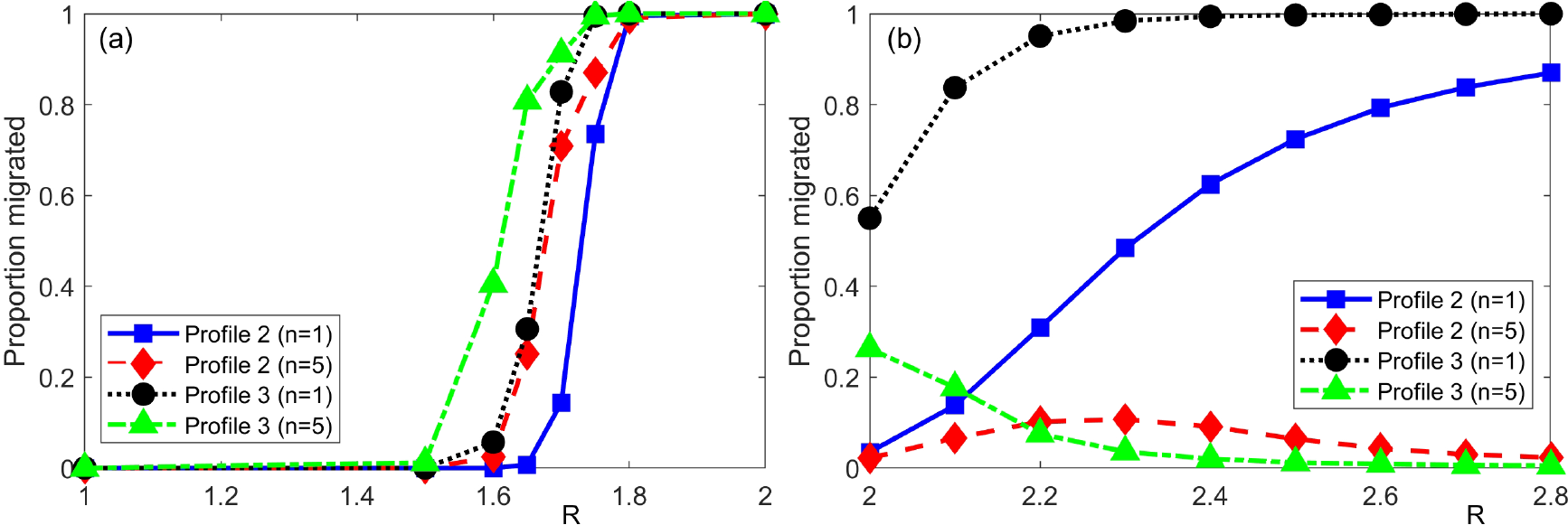
Proportion of population initially located on the lower peak that migrates to the higher peak by *t* = 5000 for various *R*, *n* and peak height ratios. (a) Results for (M3), using *κ*_3_ = 3; (b) Results for (M2), using *κ*_2_ = 0.2. Other parameters are set at Δ*y* = 0.003, *τ* = 1, *s* = 0.01.

#### 5.1.2. Influence of noisy environments

So far we have considered smoothly varying cues. In reality, environments are expected to be considerably rougher, due to small-scale variations in topography. Since a strictly local response depends on the gradient at a specific point, this variation may significantly impact on an individual’s ability to move across some environment. To explore this, we introduce a form of gradient noise into our environmental cue: we refer to Appendix C for details, but remark that the added noise contains local spatial correlation to ensure a differentiable underlying environment.

We examine the average displacement of a population moving in response to a linear topographical profile, Figure 2(d), subject to the addition of various levels of noise. Initially, the population is uniformly distributed between *y* = 0 and *y* = 1.2 (and zero elsewhere). Average displacement is calculated by numerically solving the full DAD equation (1) under twenty identically-prepared noise terms. That is, we generate twenty different noise terms according to the same process, with the only difference stemming from the numbers sampled from the (pseudo)-random number generator. We then solve (1) for each of the noise terms and calculate the overall displacement of the population, defined as the difference between the median of the population at *t* = 0 and *t* = 500. We repeat this process for different levels of noise and scale according to the average displacement under zero noise.

The results for models (M1)-(M3) are summarised in Figure 6 (a), where for the nonlocal models we consider *R* = 0.2, *R* = 0.5 and *R* = 1.0. As might be expected, increasing the strength of the noise compared to the strength of the environmental cue results in a decrease in the average displacement. Local sensing (M1) is the most susceptible to the added noise, with increasing noise significantly reducing the displacement. Nonlocal models (M2)-(M3) prove considerably more resilient, particularly for large perceptual ranges. This result is intuitive, as the nonlocal response involves averaging the environmental cue over the perceptual region, reducing the contribution of any noise. Note that as *R* → 0 for (M2), we observe convergence to (M1), consistent with our earlier findings. Overall, given that environments in nature are non-smooth and exhibit variation, nonlocal responses may be necessary for successful migration.

We now return to the example with two peaks, with two initially distinct populations, and examine how the presence of noise influences the successful migration from the lower to higher peak. We note that a small amount of noise can elicit a large change in the proportion of the population that undergoes migration (Supplementary Material, Section S2.1). To determine how different levels of variation influence hilltopping, we append twenty identically-prepared noise terms to Profile Two. Using two distinct *R* values we examine the proportion of individuals initially located on the lower peak that migrate to the higher peak by *t* = 5000, presenting the results in Figure 6 (b) for (M2) and (M3). As expected, we find that increasing the amount of noise generally reduces the ability for individuals to translocate to the higher peak for both models: for combinations of *R* and *κ* that allow the entire population to relocate in the absence of noise, introducing noise inhibits a portion of the population from doing so. Consistent with previous findings, larger perceptual ranges offset noise-induced trapping. As naturally occurring environments do not typically exhibit the smoothness of idealised models, this suggests that the mechanism by which sensory information is processed must be able to account for the noise.

**Figure 6:**
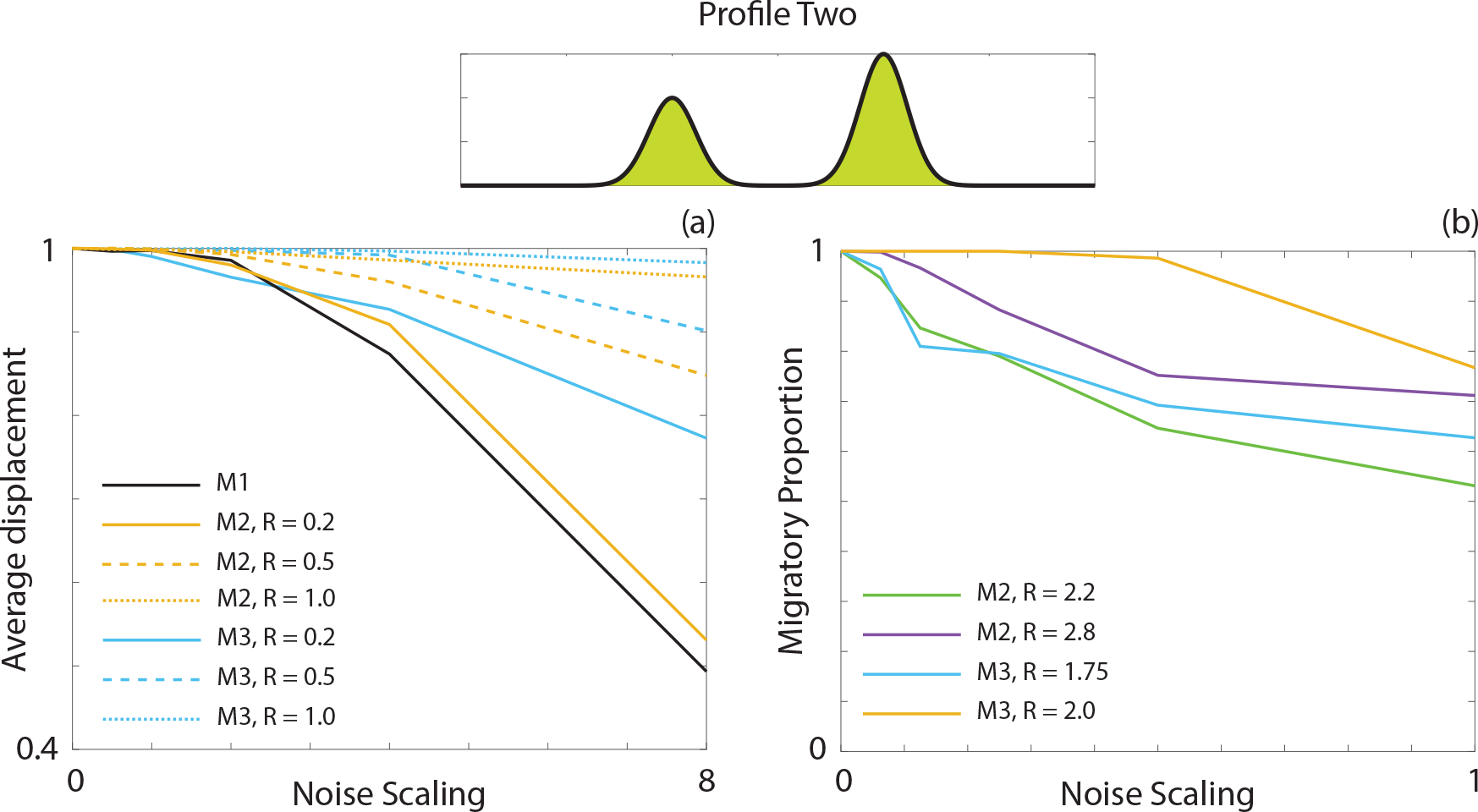
(a) Scaled average displacement by *t* = 500 for a population of individuals initially uniformly distributed between *y* = 0 and *y* = 1.2 for both local (solid) and nonlocal (dashed) responses and three *R* values. For each parameter combination the average displacement is obtained from 20 simulations on identically-generated terrain. Parameters used are *κ*_1_ = 0.3, *κ*_2_ = 0.6, *κ*_3_ = 2.0, Δ*y* = 0.003, *τ* = 1, *s* = 0.01, *n* = 1. (b) Average proportion of the population initially located on the lower peak in Profile Two that migrates to the higher peak by *t* = 5000 for M3 and *R* = 1.75 (cyan), M3 and *R* = 2.0 (orange), M2 and *R* = 2.2 (green), and M2 and *R* = 2.8 (purple). For each parameter combination the average migratory proportion is obtained from twenty simulations on identically-generated terrain. Parameters used are *κ*_2_ = 0.6, *κ*_3_ = 1.5, Δ*y* = 0.003, *τ* = 1, *s* = 0.01, *n* = 1.

**Figure 7:**
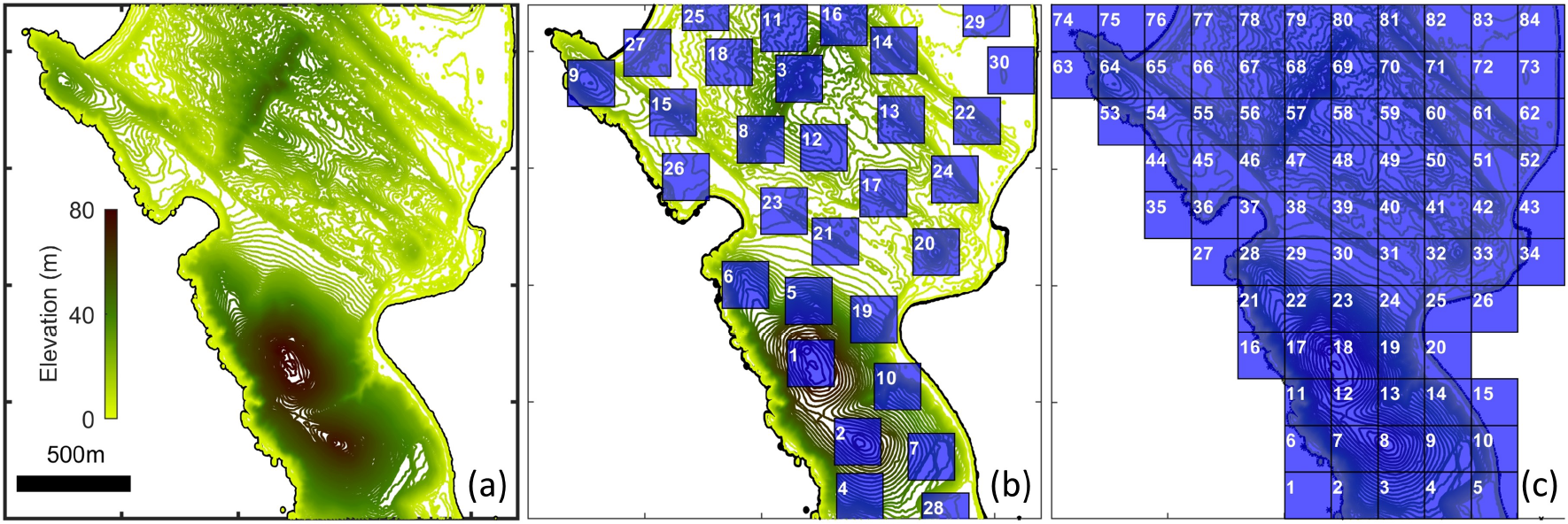
(a) Topography of Bodega Marine Reserve study site. (b)-(c) Zones used for population counts, following either (b) peak-centred or (c) tessellated gridding.

### 5.2. Bodega Marine Reserve study

#### 5.2.1. Study site and elevation data

We extend our case study to a natural environment, utilising topographical data for the Bodega Marine Reserve, a site of recent hilltopping studies [15]. High spatial resolution LIDAR data was obtained from the United State Geological Survey’s “The National Map” project over the co-ordinate range of the study site, and longitude, latitude, elevation and classification (water, ground etc.) are extracted at each data point. Interpolation onto a regular grid is performed, and the resulting topographical structure is mapped in Figure 7(a). This topographical structure allows individuals to obtain navigation information in a manner that mimics the behaviour of a population undergoing directed motion according to the topography in the Bodega Marine Reserve. A population of butterflies/moths is uniformly randomly distributed across the landmass at *t* = 0 and their movements are followed over 20 hours. Of course, more realistic distributions should account for oviposition/larval sites but a uniform distribution limits any subsequent bias in the final distributions. Simulation methods are described in the Supplementary Methods. Note that we impose boundary conditions that effectively confine the population within the bounds of the region plotted in Figure 7(a), as described in the Supplementary Methods, Section S3.

For later analyses we subdivide the land region using two methods. The first is a peak-centred scheme, Figure 7(b), where 200m by 200m (4 hectare) non-overlapping regions are defined according to topography: (i) we find the complete set of local maxima; (ii) zone 1 is centred on the highest maximum and both this maximum and any others within distance are excluded; (iii) zone 2 is centred on the next highest non-excluded maximum and so forth. The second subdivision, Figure 7(c), is a regular tessellation that intersects with the land coverage. The use of two methods is to reduce bias due to zoning.

We consider the three prototype models for topographic sensing, (M1)-(M3). Note that in order to concentrate on how altering the navigation field and its associated parameters impacts on population structuring, we fix the speed, turning rate and sensitivity coefficients at values similar to those in an earlier study [34]. Specifically, we set *s* = 4 m/min and a mean run time of *τ* = 1 min; note that these values account for pauses/resting periods during flight. Sensitivity coefficients are set at *κ*_1_ = 5, *κ*_2_ = 10 and *κ*_3_ = 0.2; the relationship between *κ*_1_ and *κ*_2_ ensures convergence from (M2) to (M1) as *R* → 0; the relationship between *κ*_2_ and *κ*_3_ is to generate quasi-equaivalent navigation fields for (M2) and (M3) when *R* = 50 m. To provide context to these values, a butterfly located on a linear ramp of gradient 10% would choose an uphill direction on just over 75% of occasions, rising to slightly more than 90% for a 20% gradient.

### 5.3. Navigation field variation

We first consider how orientation information changes as we shift from local to nonlocal models. For ease of illustration we plot only a portion of the full study site (the top left region in Figure 7(a)). Plots are presented in the form of a “direction field”, where red arrows indicate the local direction of **w** at selected points and blue lines indicate the trajectory traced out if an individual exactly follows these arrows. Light blue circles indicate trajectory end points, and can be interpreted as attractors that potentially lead to local accumulations of the population.

The plots in Figure 8 show results for (M1), i.e. *R* = 0, and (M2) with *R* = 25, *R* = 100 and *R* = 500; note that these plots assume *n* = 5. Under local sampling we observe significant fluctuations in arrow directions, with some arrows pointing in the direction of prominent peaks but others directed elsewhere. Numerous local maxima exist, so that trajectories travel short distances before becoming trapped at an attractor. Transitioning to the nonlocal model, however, dramatically reduces the number of attractors. The majority of trajectories now converge on a select few summits (examples given by black arrowheads) and trajectories can be significantly longer than the perceptual range. Effectively, movement is first according to the highest elevation angle within range, but subsequent higher peaks may be observed. For example, in Figure 8 (c) we see trajectories converging on the top-left attractor over a distance of approximately 1 km, an order of magnitude greater than the perceptual range. Increasing *R* steadily reduces the number of attractors, such that by *R* = 500 all trajectories converge on one of just two attractors within the plotted region, corresponding to *prominent peaks*. These peaks are not necessarily the two highest peaks across the studied zone, rather they correspond to the highest within their region.

Further analyses have been performed for (M2) with *n* = 1 and (M3) with both *n* = 1 and *n* = 5 (see Supplementary Materials, Section S2.2). Qualitatively we observe similar phenomena, with a reduction in the number of attractors as *R* is increased. However, a few subtleties emerge: (i) higher nonlocal orders promote the persistence of attractors at lower prominences, consistent with our earlier idealised study; (ii) for *n* = 1, attractors can occasionally form some distance away from the nearest local peak. This latter observation stems from the fact that, under a lower nonlocal order, the weighting to summit points is diluted and evaluation is more according to a broad assessment of the terrain. However, broadly the results are consistent and we note that perceptual range, rather than the precise sensing model or degree of nonlinearity, appears to be the primary determinant in the number of attractors. For the remainder of our investigation we therefore restrict to presenting the results for (M2) under *n* = 5, noting that results for the other cases are qualitatively consistent.

**Figure 8:**
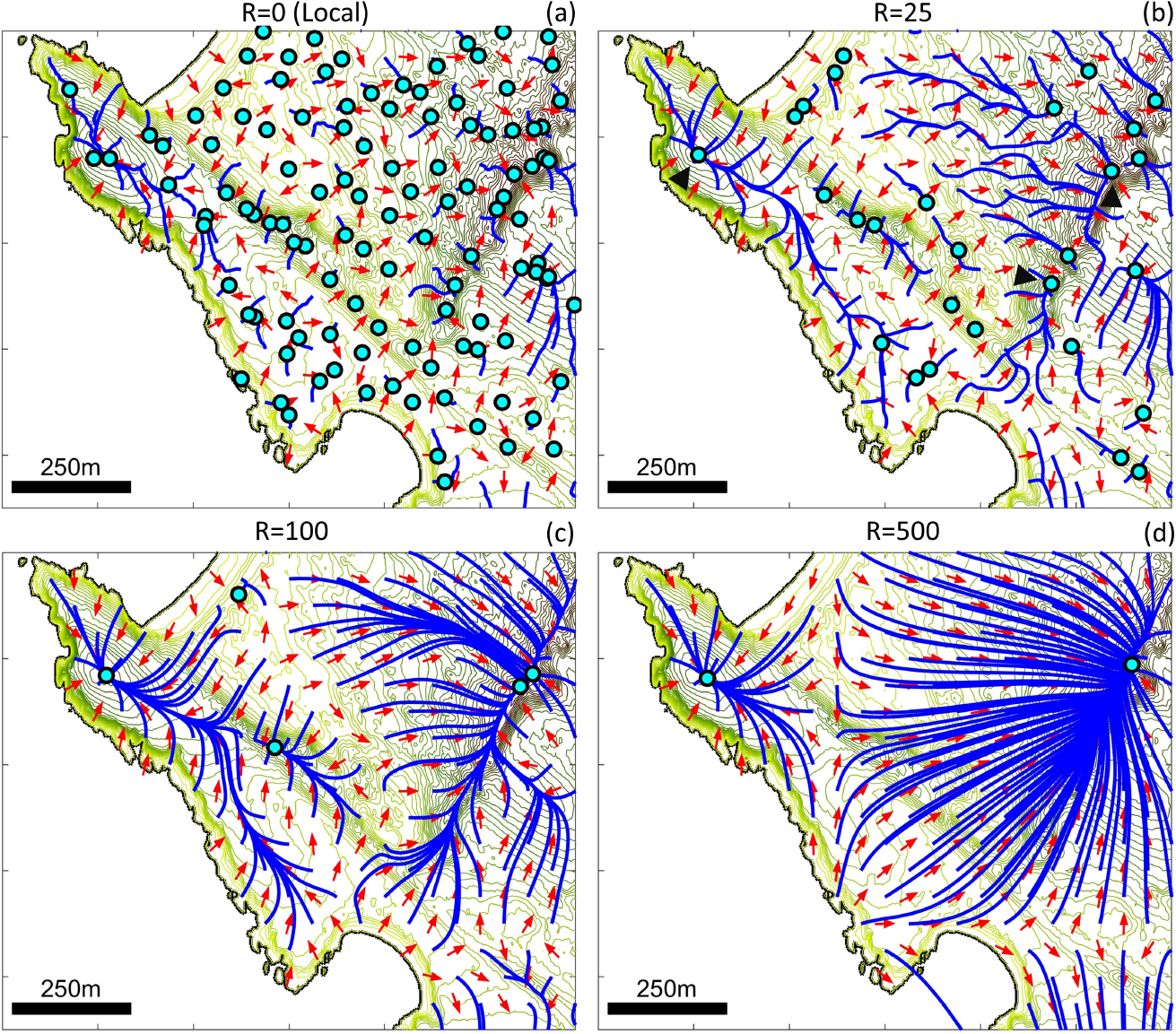
Representation of orientation information, for **w** given by (M2) with (a) *R* = 0, (b) *R* = 25, (c) *R* = 100 and (d) *R* = 500. Note that the *R* = 0 case is effectively the local gradient model (M1). For these plots we use *n* = 5 as the nonlinear coefficient.

### 5.4. Comparison of the velocity jump and macroscopic models

We proceed to simulate the VJRW and the full (unapproximated) macroscopic model (1) across a range of *R*, using (M1), i.e. *R* = 0, and (M2) with *R* = 25, *R* = 100 and *R* = 500. Results are presented in Figure 9. Agent-based model simulations of the VJRW are presented in Figures 9(a)-(d), where blue circles denote final locations (after 20 hours) and red dots indicate trajectories. The results parallel the previous direction field plots. For the local model, some larger collections occur at the most prominent peaks but the overall population remains scattered with numerous small accumulations trapped at local maxima. Expanding the perceptual range reduces trapping, allowing the population to coalesce into fewer and larger aggregates that correlate with the prominent peaks. The capacity for long-range information to influence movement is particularly illustrated by the individuals that move across the bays, attracted to peaks on the other side (arrowheads in Figure 9(d)).

Under each agent-based simulation we show the corresponding population distributions predicted by the macroscopic model after 2, 8 and 20 hours. Note that the distributions correlate well with those of the agent-based model, suggesting that the macroscopic model accurately captures individual-level behaviour at the spatial and temporal scales of study. We subsequently focus on the macroscopic model, exploiting its computational advantages. The simulations confirm the results of the agent-based model, with large sampling radii allowing the population to coalesce into fewer and larger aggregates: for example, for *R* = 500 we observe the population has coalesced into just 3 principle populations.

We also test the validity of using an approximation, comparing the full DAD model with its first order approximation (assuming a weak cue) in Figure 10. Subtle difference emerge in regions corresponding to the strongest field strengths but, overall, the first order approximation performs remarkably well: differences between the approximation and full DAD model are likely to be small in comparison to errors arising from parameter estimation. Moreover, differences are primarily noticeable at early times, with the results in the quasi-steady state distribution, presented in Figures 10(c)-(d), visually indistinguishable.

**Figure 9:**
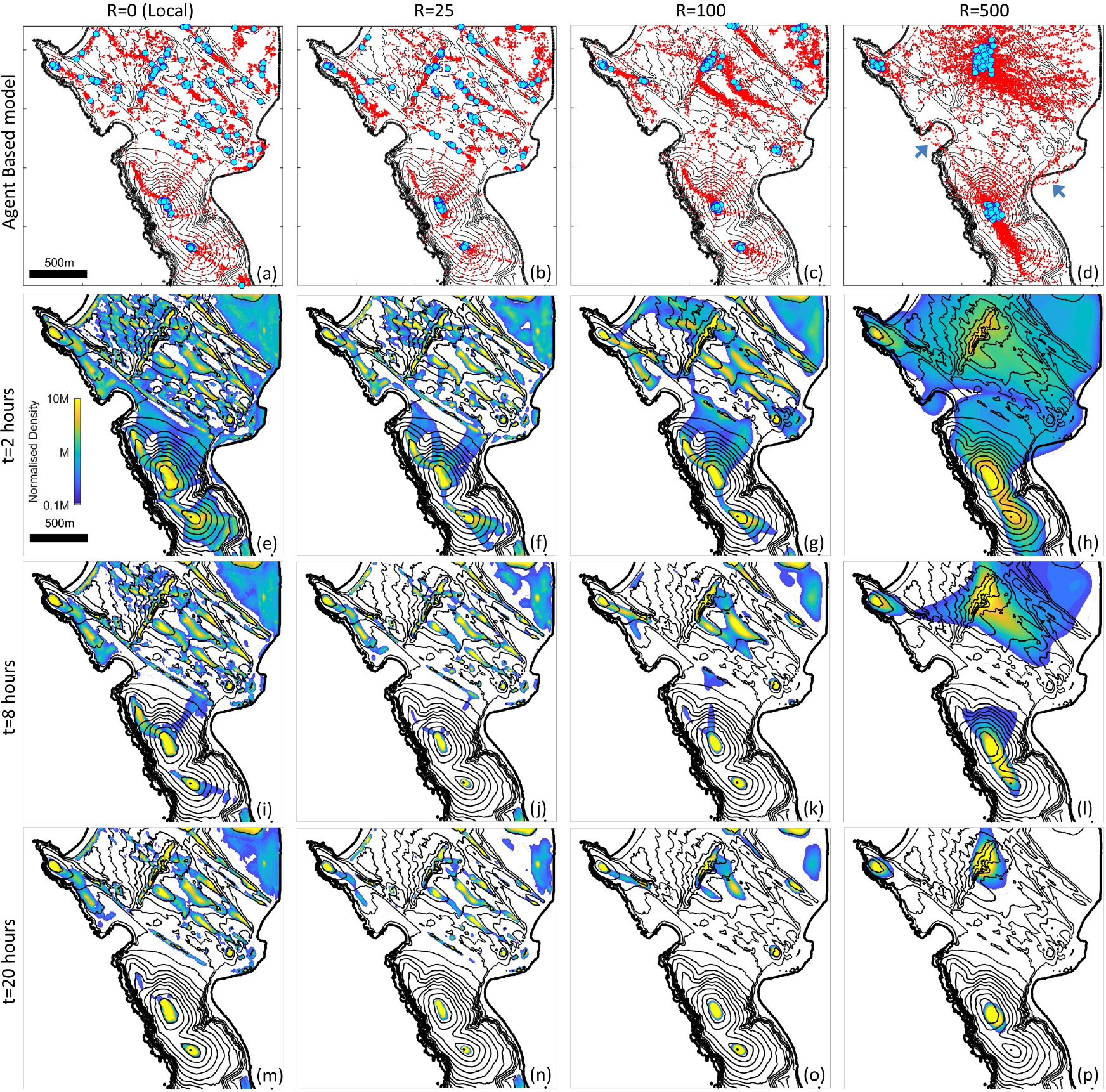
Simulations of the VJRW and full DAD model (1) using elevation data for the Bodega Marine Reserve. Navigation field given by (left to right) the local model (M1), i.e. *R* = 0, and (M2) with *n* = 5 and *R* = 25, *R* = 100 and *R* = 500. (a)-(d) Simulations of the VJRW. Blue circles represent end locations and red dots indicate trajectories, for 250 individuals that are initially randomly uniformly distribued across the landmass. (e)-(p). Corresponding population distribution predicted by (1) at (e)-(h) 2, (i)-(l) 8 and (m)-(p) 20 hours. Density plotted according to the colourbar in (e), where *M* denotes the the density if the population is uniformly distributed across the landmass. Parameter values are as described in the main text.

**Figure 10:**
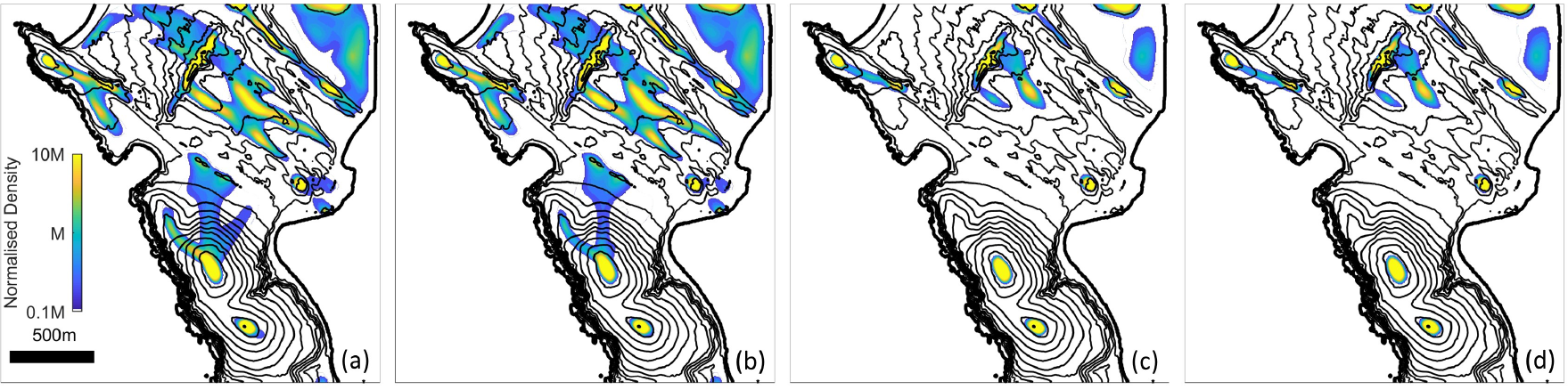
Comparison between the full DAD model (1), (a) and (c), and the first order approximation obtained under the assumption of a weak cue, (b) and (d): (a,b) *t* = 4 hours; (c,d) *t* = 20 hours. Simulation details as in Figure 9, where for this comparison we use (M2) with *n* = 5, *κ*_2_ = 10 and *R* = 100.

We perform an analysis of coalescence by counting population densities present in different zones, under the two schemes described previously. Results are consistent regardless of zoning scheme, and we therefore only present the data for the peak-centred scheme. In Figure 11(a) we present bar charts, which display the number of zones (at 20 hours) where the zone population density is at 2×, 5×, 10 and ×20 the mean density level *M*, defined as the density if the population is uniformly distributed across the landmass. Local/short range sampling generates numerous sites with moderately raised densities, but very few with significantly increased densities. For abundant populations these moderate increases may offer mating benefits, but for rare species they may have minimal effect on the overall encounter rate. Longer range sampling, however, generates relatively few sites with raised densities, but those sites exhibit substantial increases in population density. The conferred advantage is likely to be particularly acute for the rarest populations, substantially raising densities and increasing the likelihood of encounters.

We further calculate the *normalised encounter rate*, which we define as

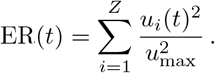

In the above, *Z* is the total number of zones, *υ*_*i*_(*t*) is the density of individuals within zone *i* at time *t* and *u*_*max*_ is the maximum zone density (i.e. the density if the entire population is concentrated into a single zone). Note that the above assumes that male/female distributions are equivalent and that encounter rates are proportional to the product of male and female densities. The resulting value ranges between 0 and 1. Plotted as a function of time, nuances are revealed in terms of how the perceptual range impacts on the encounter rate. Over short time ranges lower to medium values of *R* appears to be more advantageous; at longer times, larger *R* values are optimal. Intuitively, larger perceptual range can encourage direct movements towards distant summits and less frequent on route encounters; lower to medium *R* may facilitate earlier encounters. This is reflected by comparing the trajectories indicated by Figure 9(c) and (d). For *R* = 100 we observe trajectories (indicated by red dots) densely channelled along “virtual corridors” [41], which subsequently become dispersed for *R* = 500. Given the short lifespans of many insects, and the potential need to mate rapidly, sampling across a very large portion of the environment may therefore not always be the optimal strategy.

**Figure 11:**
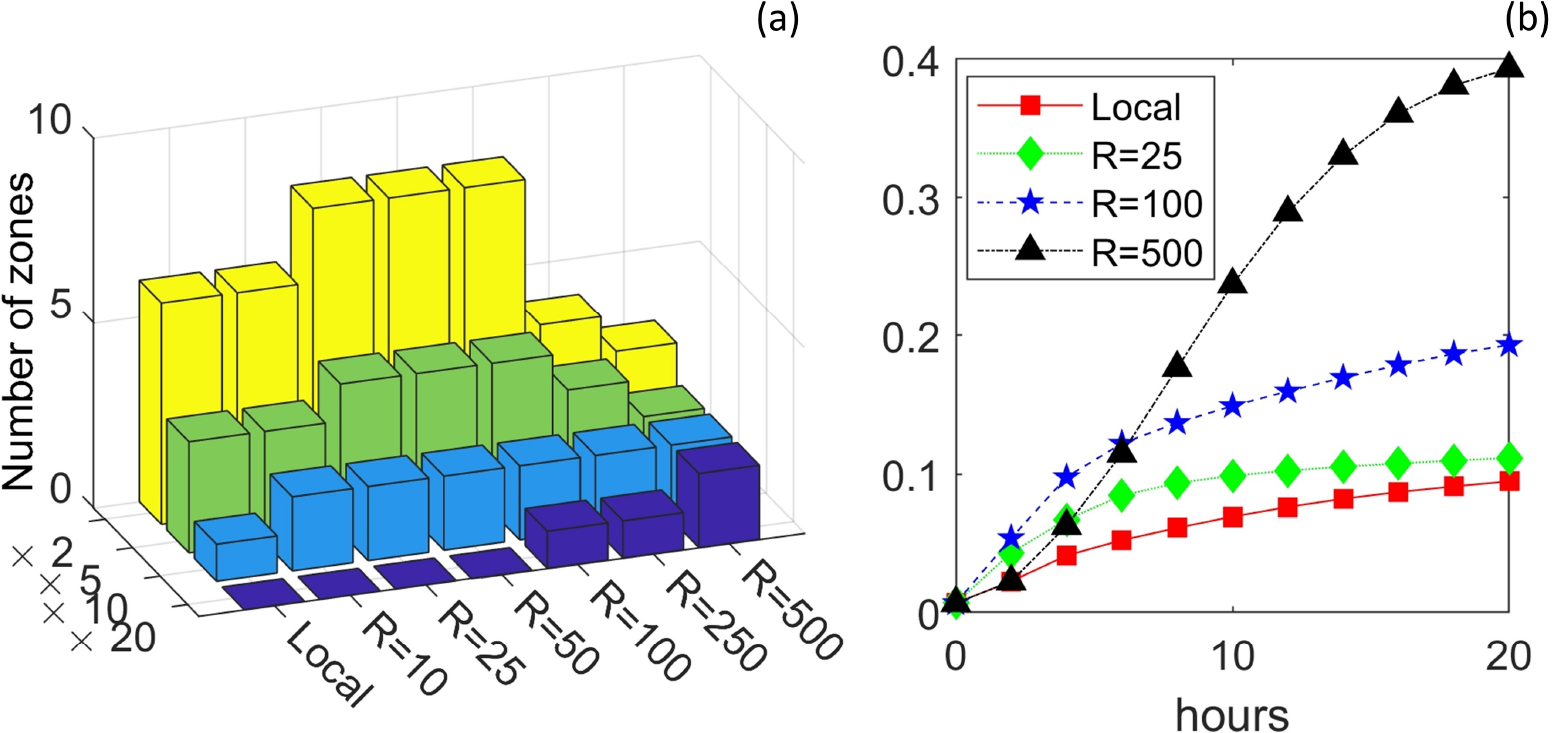
Results of zoning analysis for (M1) and (M2) under various *R*. (a) At the end of the simulation (*t* = 20 hours) we calculate the number of zones for which the population has increased to 2×, 5, 10, 20 the mean density.(b) We plot the normalised encounter rate as a function of time for (M1) and (M2) under various *R*. All results use the peak-centred zoning method while model parameters are as described in the text.

## 6. Discussion and conclusions

The ability to detect information located distant to a cell or organism can be crucial for its navigation and movement. Larger organisms can achieve this through various means: vision and hearing provide obvious, pervasive examples; electrolocation by fish through their lateral line organs presents a more specialised case [20]. Certain migrating cells can extend long protrusions (filopodia, cytonemes) that span multiple cell diameters [50]. The majority of theoretical models, however, implicitly assume that the underlying cue is local (or effectively local) with respect to the individual’s position: for example, a chemical gradient computed across the individual’s basic body dimension. Here we have described a multiscale framework, based on a biased velocity-jump random walk, where the orienting environmental cue is either local or distant to the individual’s current location.

Our underlying random walk is standard and, when scales of interest are suitably macroscopic, its behaviour can be described by a general form nonlinear drift-anisotropic diffusion equation [18], the parameters of which depend on the navigation field/environmental cue. Via standard expansions of the modified Bessel functions, we show that in certain parameter regimes we can approximate the general form equation as a Keller-Segel type equation (for local gradient sensing) and a nonlocal integro-partial differential equation (under nonlocal sensing). Nonlocal integro-PDEs have become popular in movement models, in both ecological [28, 7, 10] and cellular systems [33, 17, 36]. The work here provides a means of connecting these models with microscopic/individual level behaviour; for other recent approaches see [27, 4]. Of course, the form of continuous model is intrinsically linked to the VJRW assumptions and their alteration could demand recalculation. For example, the default assumption of exponentially distributed runtimes does not universally apply: under certain scenarios, occasional long transits are observed, a so-called Levy process described by long-tailed power-law distributions. Generalising VJRWs to include other runtime distributions or resting periods between runs has been considered in a number of studies (e.g. see [16, 49, 9] amongst others). Even more challenging is the assumption of stochastically-independent walkers: while reasonable for scattered/dispersed population, this becomes questionable if the individuals aggregate/cluster tightly.

Explicit descriptions of (ecological) perceptual range have been included in a variety of theoretical/computational models, including agent-based [40], stochastic mechanistic resource selection models [2] and integro-PDEs of drift/diffusion type [10]. Our modelling here has similarities, in particular, to the latter two studies. Similar to [2], our modelling is founded on stochastic random walk movements of an individual evaluating landscape across a perceptual range; in [29] the authors similarly derive diffusion-advection approximations for their stochastic model. The integro-PDE model studied in [10] includes a drift component that features nonlocal gradient following to some resource, similar to the form (11) suggested as a first order approximation to our general model.

The utility of the model was demonstrated through a specific application to hilltopping behaviour. Under both idealised and genuine terrain profiles we demonstrated that nonlocal sensing allows a population to overcome terrain noise, so that individuals transit from lower to higher peaks; local sensing, on the other hand, can lead to trapping of population members at local maxima close to their starting location. Since topographical data will typically possess numerous local maxima, this suggests that an effective hilltopping mechanism would be based on highly nonlocal sensing. Applied to the topographical data of the Bodega Marine Reserve, nonlocal sensing substantially increases population densities at a subset of “prominent peaks” across the studied area. As such, the model broadly reproduces field findings on the distribution of tiger moths (*Arctia virginalis*) at the same site [15]. A direct translation between our results and the field findings of [15] would require additional refinements, for example accounting for the biases arising from larval site/vegetation distribution. Here we have not accounted for this, allowing us to focus exclusively on the role played by nonlocal assessment on population distribution. Perceptual range, rather than the exact mode of sensing or nonlinear order, appears to be the primary determinant in the number of congregation sites. Large perception ranges are likely to be particularly helpful for the scarcest populations, bringing the dispersed population into concentrations at a small number of sites. Yet, the advantages become less clear-cut at shorter timescales, where the more dispersed travel routes taken to reach distant sites may reduce the likelihood of earlier mating: effectively, the narrow “virtual corridors” [41] encouraged by lower perceptual ranges become dispersed for larger *R*. Of course, the present modelling neglects to account for adaptive strategies, where individuals may first preferentially move to a nearby (lower) maximum and subsequently move to one more distant if no encounters are formed. Such hypotheses can easily be tested through more sophisticated nonlocal sensing rules.

Here we consider two prototype models for nonlocal sensing: a nonlinear gradient and a non-linear maximum model. In the context of hilltopping, these have clear and direct interpretations regarding whether the bias is according to the highest elevation angle or elevation within perceptual range. Either model appears to be an effective mechanism for overcoming terrain roughness and generating fewer, higher populated accumulations, although we remark that our earlier idealised experiments suggest that the nonlinear maximum model may be slightly more effective for overcoming noise. From a human perspective, elevation angle is easily gauged but estimating true height would require an additional capacity for depth perception. Overall, though, these prototypes can serve as simple models for describing various forms of sensing, such as visual, auditory, or chemical information. However, focused applications would clearly demand closer inspection of their foundation. For example, the choice of uniform Ω is somewhat simplistic, where we may expect a more gradual drop-off in the ability to detect at a distance. Clearly the more general form (7) allows for a lot of tailoring.

The hilltopping application features a fixed environmental cue, however, often the cue will vary with time. This could occur independently of the population, such an odour transported by turbulent currents, or according to the population distribution, such as a cell population that internalises a chemical signal or whales communicating through song. A key study would be to extend the analysis here to examine how nonlocal sensing impacts on oriented movements under dynamic variation of the cue; steps in this direction have been made in [10], where the potential beneficial effects of nonlocal sensing were illustrated. Related to this, here we have restricted ourselves to consider only fixed noise. Time-varying noises in the cue, such as molecular fluctuations in a chemical signal, are of significant interest: fluctuations may further restrict the effectiveness of local gradient-based sensing mechanisms and highlight the importance of nonlocal sensing. Studies of this nature may provide insight into some key questions, including the impact of human activity on ecological populations. For example, the effective perceptual range of cetacean auditory communication has been heavily eroded by noise pollution, such as commercial fishing, naval sonars and oil/gas exploration [38, 53]. Detailed modelling of such problems via our framework may provide important predictions on how such unwanted noise can impact on population dynamics.

## Acknowledgements

STJ thanks the Australian Mathematics Society for the award of a Lift-Off Fellowship to visit KJP. KJP acknowledges the Dipartimento di Scienze Matematiche and the Politecnico di Torino for a Visiting Professorship award.

**Figure A.1.**
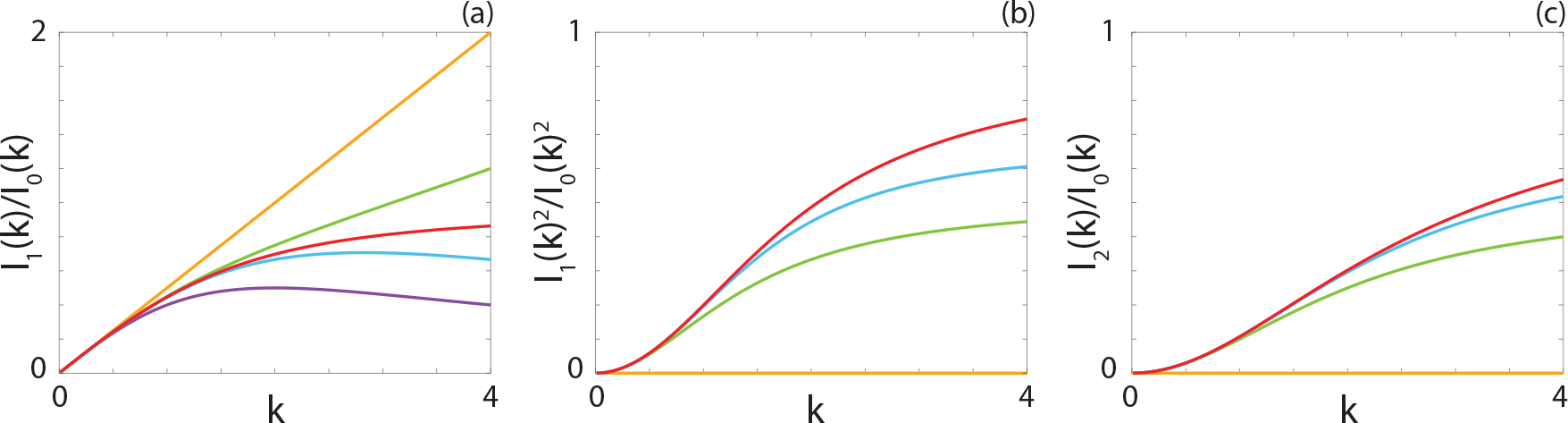
Comparison between the ratio of modified Bessel functions (red) (a) *I*_1_(*k*)/*I*_0_(*k*), 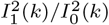, (c) *I*_2_(*k*)/*I*_0_(*k*), and the *O*(*k*) (orange), *O*(*k*^2^) (purple), *O*(*k*^3^) (green) and *O*(*k*^4^) (blue) approximations of the ratios. Note that the *O*(*k*^2^) and *O*(*k*^3^) approximations are the same for 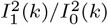 and *I*_2_(*k*)/*I*_0_(*k*).

## Appendix A Modified Bessel function expansions

## Appendix A.1. Small argument expansions

Under nonnegative integer orders, modified Bessel functions of the first kind have the following power series expansions [31]

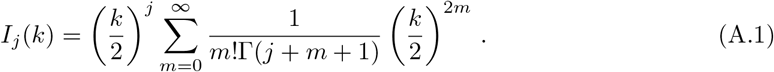

Given (5) and (6), we are particularly interested in *j* = 0, 1, 2 and the ratios *I*_1_(*k*)*I*_0_(*k*), 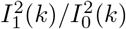 and *I*_2_(*k*)/*I*_0_(*k*). Using (A.1), we first approximate the modified Bessel functions where *j* = 0, 1, 2 as

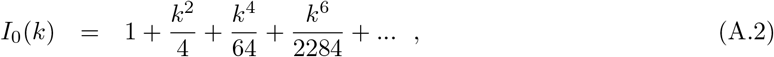

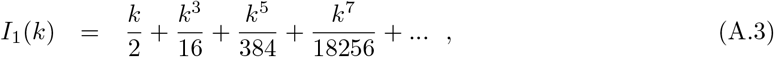

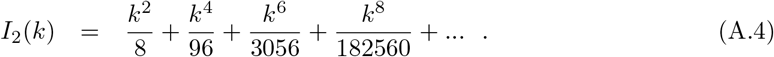

To obtain the relevant ratios, *I*_1_(*k*)/*I*_0_(*k*), 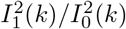 and *I*_2_(*k*)/*I*_0_(*k*), we truncate the polyno-mials on the numerator and denominator such that terms up to *O*(*k*_*n*_) are included. We obtain

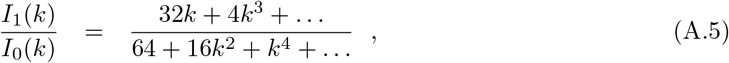

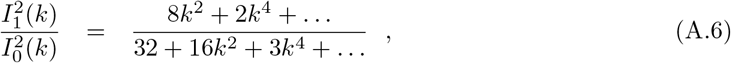

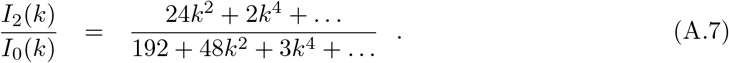

In general, according to the size of *k* we can truncate the above calculations at the *k*^*n*^th term to obtain a reasonable approximation: Figure A.1 presents a comparison between a numerical approximation of the relevant ratios modified Bessel functions and the *O*(*k*^*n*^) approximation of each of the ratios. For *k* < 2 the ratios are adequately approximated by at least one of the truncated power series and, as expected, including additional terms results in a better approximation. As *k* increases the distance between the approximations are less accurate, suggesting that it would be inappropriate to implement these approximations in (1) above some threshold *k* value.

**Figure A.2.**
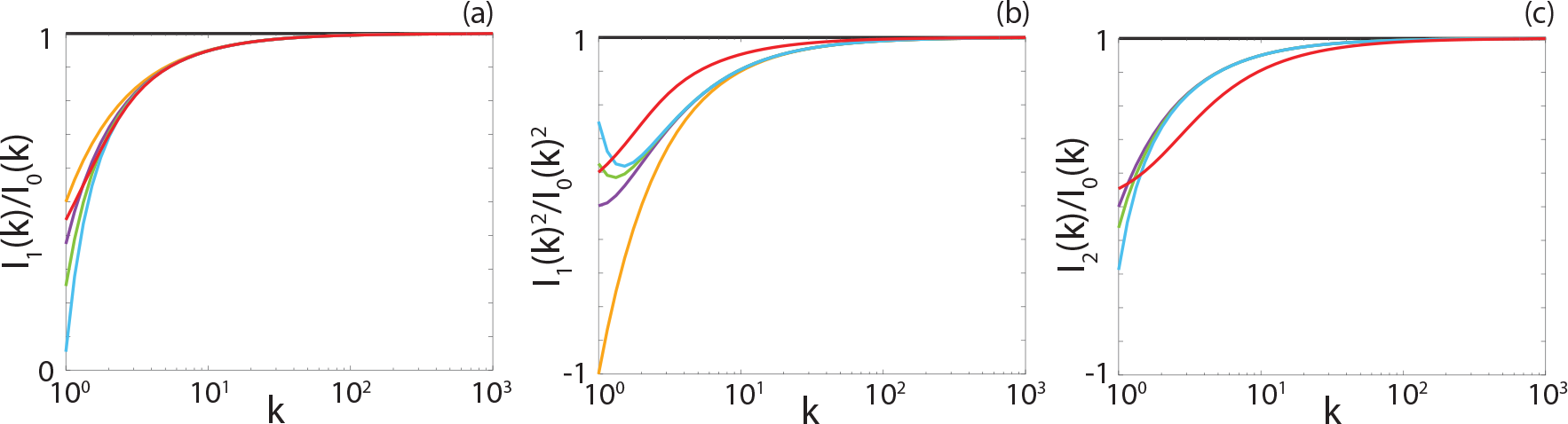
Comparison between the ratio of modified Bessel functions (red) (a) *I*_1_(*k*)/*I*_0_(*k*), 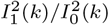, (c) *I*_2_(*k*)/*I*_0_(*k*), and the *O*(1), *O*(*k*^−1^) (orange), *O*(*k*^−2^) (purple), *O*(*k*^−3^) (green) and *O*(*k*^-4^) (blue) approximations of the ratios. Note that the *O*(*k*^−1^) and *O*(*k*^−2^) approximations are the same for 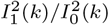.

## Appendix A.2. Large argument approximations

For larger *k* we can instead exploit the following large argument approximations for the modified Bessel functions of first kind [31], where for order *j* we have

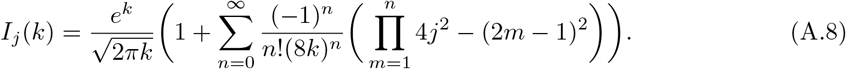

Again, we are primarily interested in ratios *I*_1_(*k*)/*I*_0_(*k*), *I*_1_^2^(*k*)/*I*_0_^2^(*k*) and *I*_2_(*k*)/*I*_0_(*k*). As the summation component of (A.8) is less than one for large *k*, we can approximate the relevant ratios as

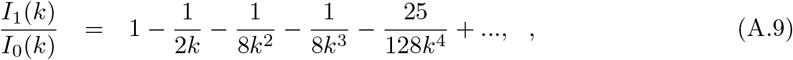

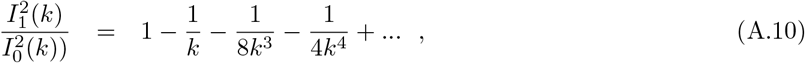

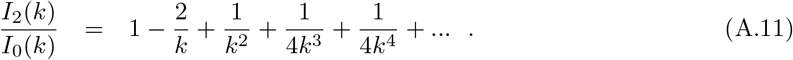

As A comparison between the approximations truncated at *O*(*k*^*n*^) for *n* = 0, 1,…, −4 and the numerical approximation of each of the ratios of modified Bessel functions is presented in Figure A.2. We observe that all approximations for *I*_1_(*k*)/*I*_0_(*k*) are indistinguishable from the numerical approximation for *k* > 10^1^, whereas for both 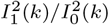 and *I*_2_(*k*)/*I*_0_(*k*) the approximations are near exact if *k* < 10^2^. Interestingly, the *O*(*k*^−1^)−*O*(*k*^−4^) converge to each other before converging to the numerical solution, which suggests that there is limited benefit to higher order approximations for large *k* values.

## Appendix B. Equation approximations

Here we only state approximations for model (M1); similar approximations can be formulated for (M2)-(M3) following an analogous process. Note that we suppress subscripts on the *κ* sensitivity coefficient for clarity of presentation.

## Appendix B.1. (M1) under small arguments

Assuming small *k* (shallow gradient ∇*E*/small response coefficient *κ*), we approximate (5)-(6) using the first few terms of the expansions (A.5)-(A.7). The series of *O*(*k*), *O*(*k*^2^), *O*(*k*^3^),…expansions are as follows:

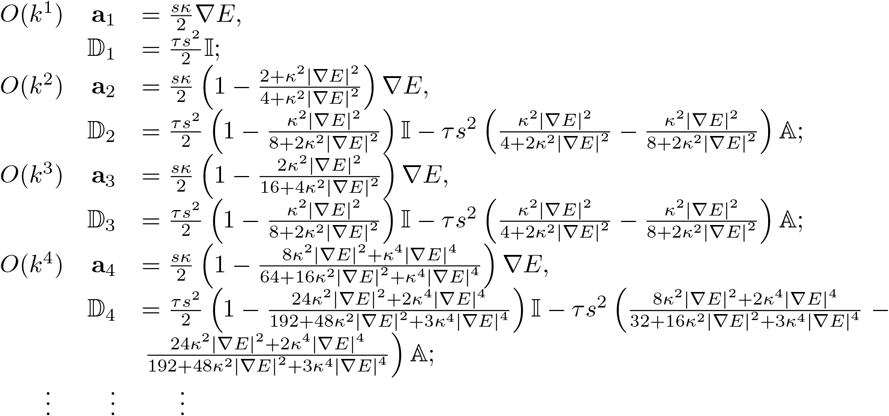

## Appendix B.2. (M1) under large arguments

For large arguments *k* (shallow gradient ∇*E*/small response coefficient *κ*), we can utilise the first few terms of the expansions (A.9)-(A.11). Substituting into (56) generates a sequence of *O*(*k*^0^), *O*(*k*^−1^), *O*(*k*^−2^),… expansions as follows:

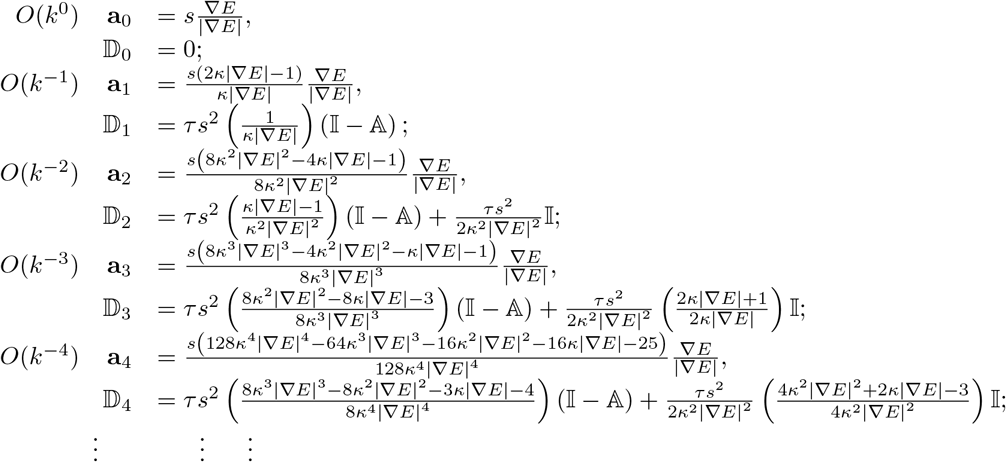

## Appendix C. Terrain noise

To generate smoothly varying noisy terrain we consider a form of fractal noise [44]. We first sample numbers from the normal distribution with mean zero and standard deviation one at *m* points. We interpolate these numbers onto a grid with 2(*m* - 1) + 1 points using spline interpolation, and take the first *m* points from the grid. This ensures that any two noise points are correlated. We repeat the process, generating *n* random values and interpolate these values onto a grid with 4(*m* - 1) + 1 points, taking the first *m* points from this new grid, and adding this noise to the Previously generated noise. This process is repeated log_2_(*y*/(2^*f*^ Δ*y*)) times. Here we take *f* = 3. Finally we apply a three point hat filter to smooth the noise further and scale it such that it lies between −0.5 and 0.5.

## Supplementary Materials

### S1. Supplementary results for approximations

In Figure S.1 we present a comparison between the full DAD model solution and the *O*(*k*) to *O*(*k*^4^) approximations for three *s* values. As stated in the main manuscript, the approximations are independent of *s* and hence the disparity between the average behaviour of the VJRW and the continuum approximations are not due to the approximations of the Bessel functions.

### S2. Supplementary results for hilltopping study

#### S2.1. Idealised experiments

We compare model (M1) and (M3) to determine their suitability for hilltopping. Individuals are initially uniformly distributed on 0 ≤ *y* ≤ 2. The solution profiles for the average behaviour of the VJRW, full DAD model solution and *O*(*k*) to *O*(*k*^4^) approximations are presented in Figure S.2. We observe that by *t* = 10, for both the local and nonlocal models, the number of individuals located on the peak has increased. As time increases, more individuals migrate onto the peak such that by *t* = 100 the majority of the population is located on the peak. Notably, the population undergoing a nonlocal response are more dispersed across the peak. Furthermore, less of the non-local population is left near domain boundaries. This is intuitive, as the nonlocal nature of the response means that individuals are able to detect the presence of the peak from a distance. In contrast, the local gradient is close to zero and hence the movement of individuals is dominated by random motility. Similar to the previous results, the macroscopic PDE description of the population density accurately matches the average velocity-jump behaviour. Additionally, the *O*(*k*) to *O*(*k*^4^) truncations approximate the full DAD model results well for both the local and nonlocal responses, particularly as *t* increases.

**Figure S.1:**
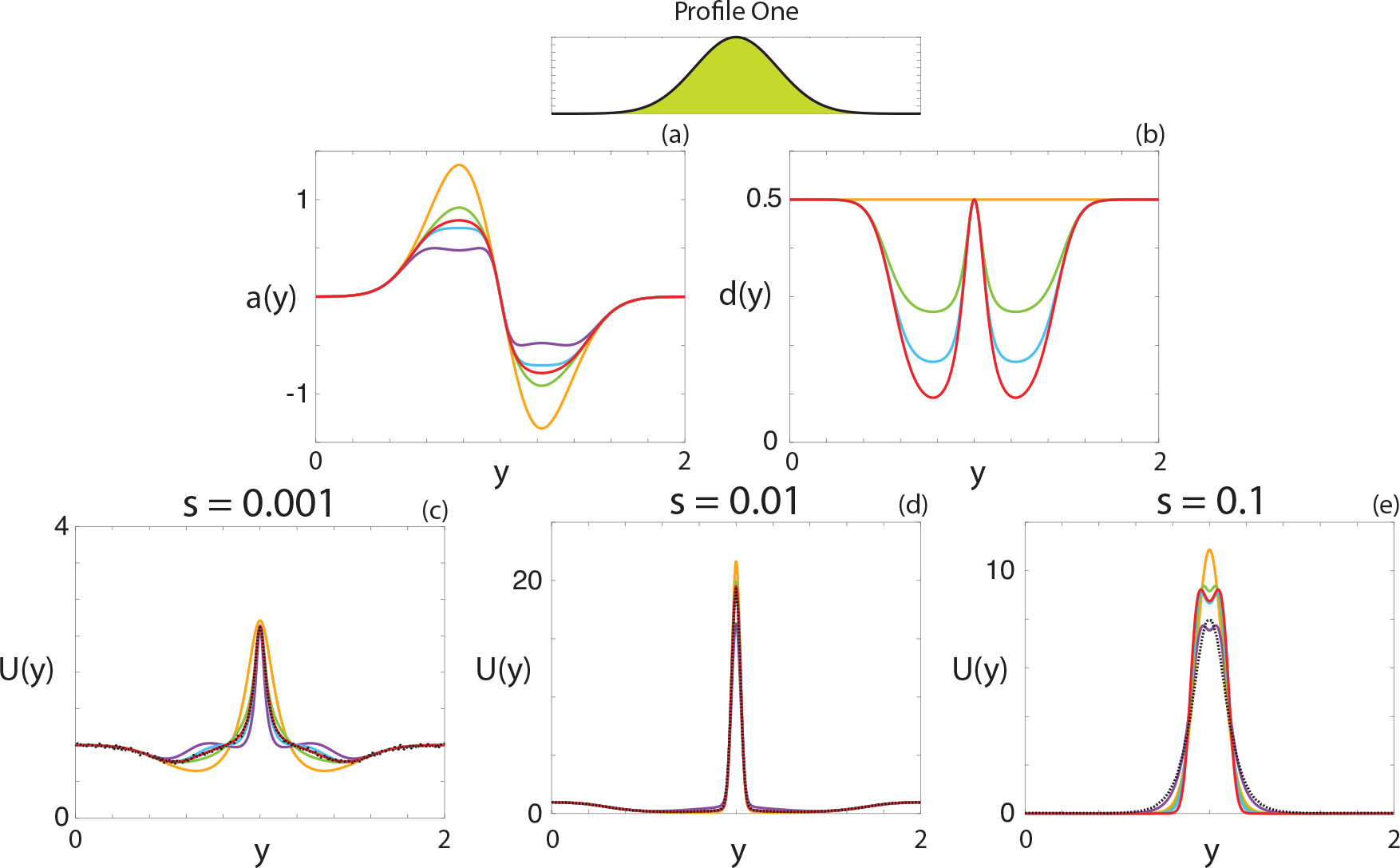
(a) Comparison between the drift-velocity (red), and the corresponding *O*(*k*) to *O*(*k*^4^) approximations (orange, purple, green, blue, respectively). (b) Comparison between the diffusivity function (red), and the corresponding *O*(*k*) to *O*(*k*^4^) approximations (orange, purple, green, blue, respectively). (c)-(e) Comparison of the average velocity-jump behaviour (black, dashed), and the numerical solutions to the full DAD model (red) and the *O*(*k*) to *O*(*k*^4^) approximations (orange, purple, green, blue, respectively) for Profile One for (c) *s* = 0.001, (d) *s* = 0.01, (e) *s* = 0.1. Parameters used are *t*_end_ = 100, *β* = 10, *κ*_1_ = 1, *M* = 10^6^, Δ*y* = 0.001.

**Figure S2:**
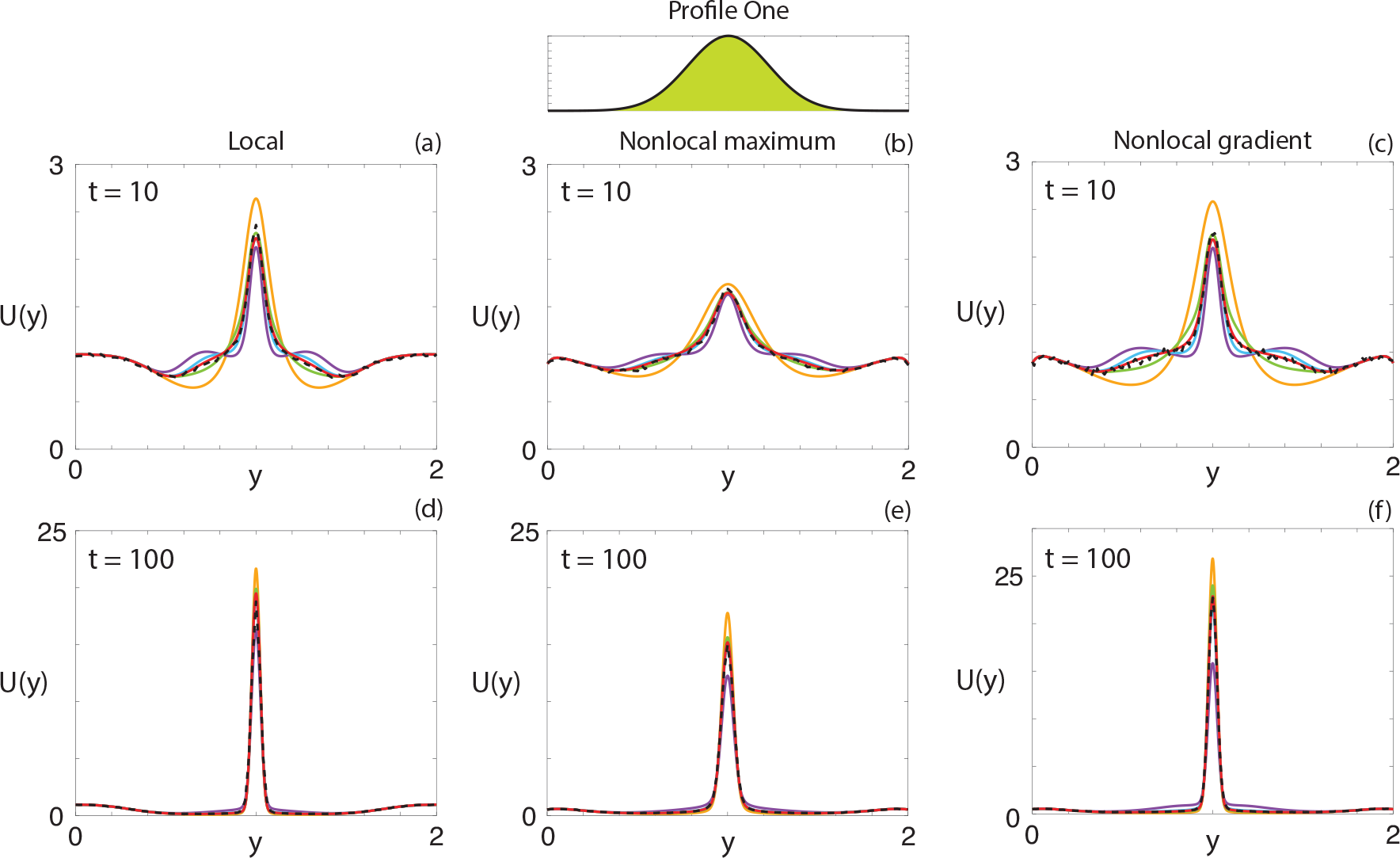
Comparison of nonlocal and local sensing mechanisms for Pro_le One. Individuals are initially uniformly distributed. Parameters used are *κ*_1_ = *κ*_3_ = 3, *κ*_2_ = 0.3, *M* = 2 × 10^5^, Δ*y* = 0:001, τ = 1, *s* = 0:01

To investigate the influence of the nonlinear order and the perceptual range we consider a population of individuals that are uniformly distributed on 0 ≤ *y* ≤ 6, with an environmental cue corresponding to Profile Two for both the nonlocal maximum and nonlocal gradient models, and analyse the distribution of individuals at *t* = 500. The solution profiles for *R* = 0.1, 0.5, 1.0, 1.5 and *n* = 1, 5 are presented in Figure S.3. For all cases we observe that there are more individuals located on the higher of the two peaks. With the exception of *R* = 1.5 for the nonlocal maximum model, changing the nonlinear order does not significantly change the solution profiles for these parameters. As discussed in the main manuscript, this is due to lack of change in the dominant direction between *n* = 1 and *n* = 5 for these parameter combinations.

We now consider the case where the environmental cue corresponds to Profile Two but with two initially separated populations located on the two peaks for *R* = 1.5 and *R* = 2.0, and present the results in Figure S.4. We see that for *R* = 1.5 the two populations remain distinct for both *n* = 1 and *n* = 5. Increasing the perceptual range such that *R* = 2.0, we observe that the two populations begin to mix. Interestingly, this only occurs on the higher peak, as only the population initially located on the lower peak undergoes migration with the population initially located on the higher peak preferring to remain. For *n* = 5 and *R* = 2.0 we observe that the population on the higher peak is more concentrated than for other parameter combinations.

We again consider the example with two peaks, with two initially separated populations, and examine how the presence of noise influences the successful migration from the lower peak to the higher peak. In Figure S.5 we present a comparison between two environments generated in an identical manner - the same parameters are used, and any difference is due to the random nature of the noise. We note that the deterministic component of this environment is the same as Profile Two, detailed previously. Interestingly, we see a dramatic difference in the proportion of the population initially located on the lower peak that migrates to the higher peak during the simulation. For the environment presented in Figure S.5(a), the entire population initially located on the lower peak remains on that peak throughout the simulation, whereas for the environment presented in Figure S.5(a), the majority of the population initially located on the lower peak migrates to the peak. If we compare the height of the terrain either side of the lower peak, we observe that the height in the negative y-direction is, on average, higher than the height in the positive y-direction (excluding the higher peak). As such, a nonlocal response that depends on the terrain height across a perceptual range will be more likely to move in the negative y-direction in this case, which hinders migration to the higher hilltop. We re-iterate that all parameters are the same in the simulations corresponding to Figures S.5(c)-(d), and that the only difference is due to the randomly generated noise. The difference in the observed migration between the two environments highlights the potential for small-scale variation to influence hilltopping, even in the presence of a nonlocal sensing mechanism.

**Figure S.3:**
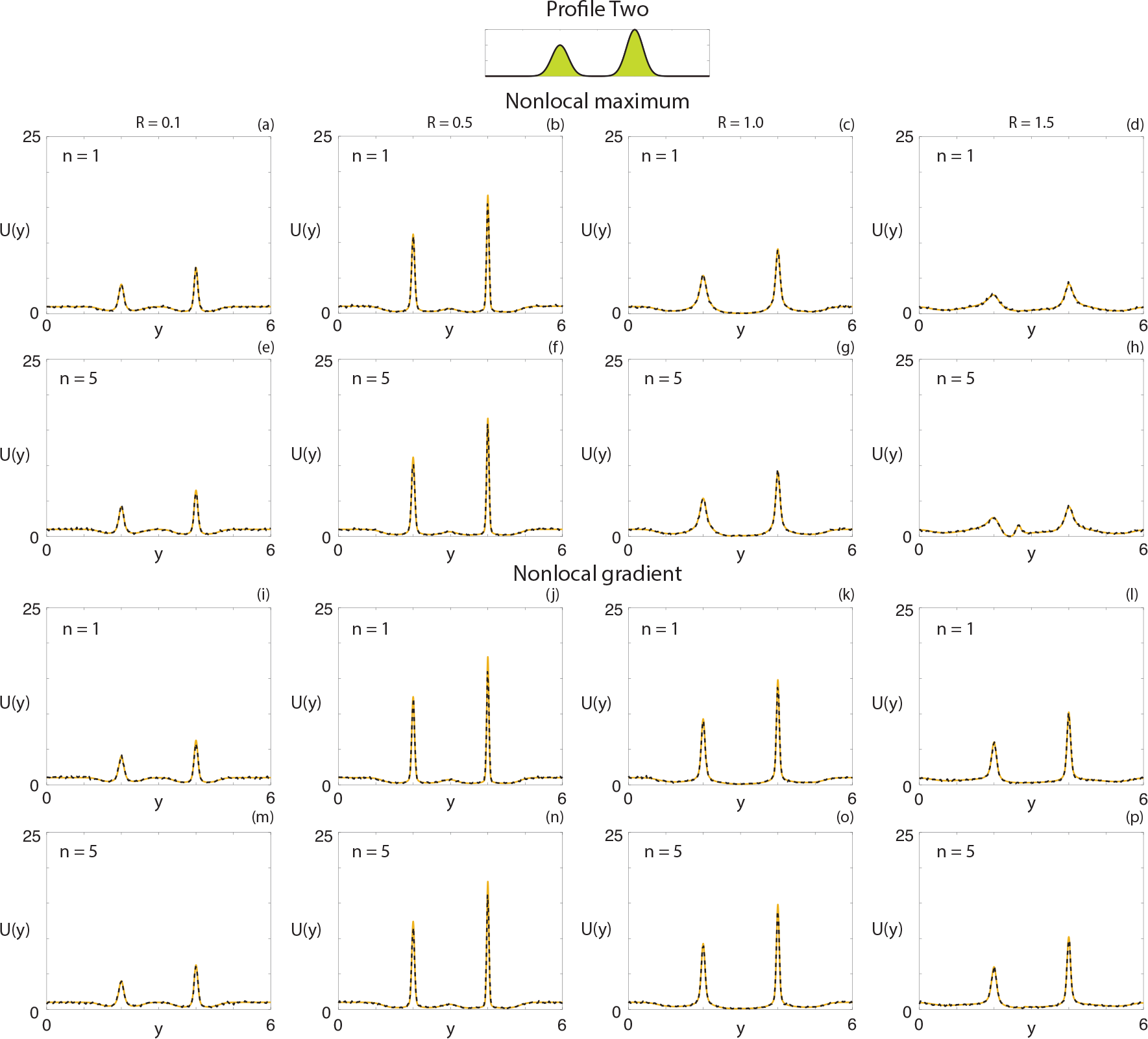
Comparison of different sensing radii and nonlinear order for the full DAD model (orange solid) and the average velocity-jump process (black dashed), respectively, using (a)-(h) the nonlocal maximum model and (i)-(p) the nonlocal gradient model. Individuals are initially uniformly distributed. The environmental cue corresponds to Profile Two. Parameters used are *t*_end_ = 1000, (a)-(h) *κ*_3_ = 3, (i)-(p) *κ*_2_ = 0.3, *M* = 2 × 10^5^, Δ*y* = 0.003, *τ* = 1, *s* = 0.01.

**Figure S4:**
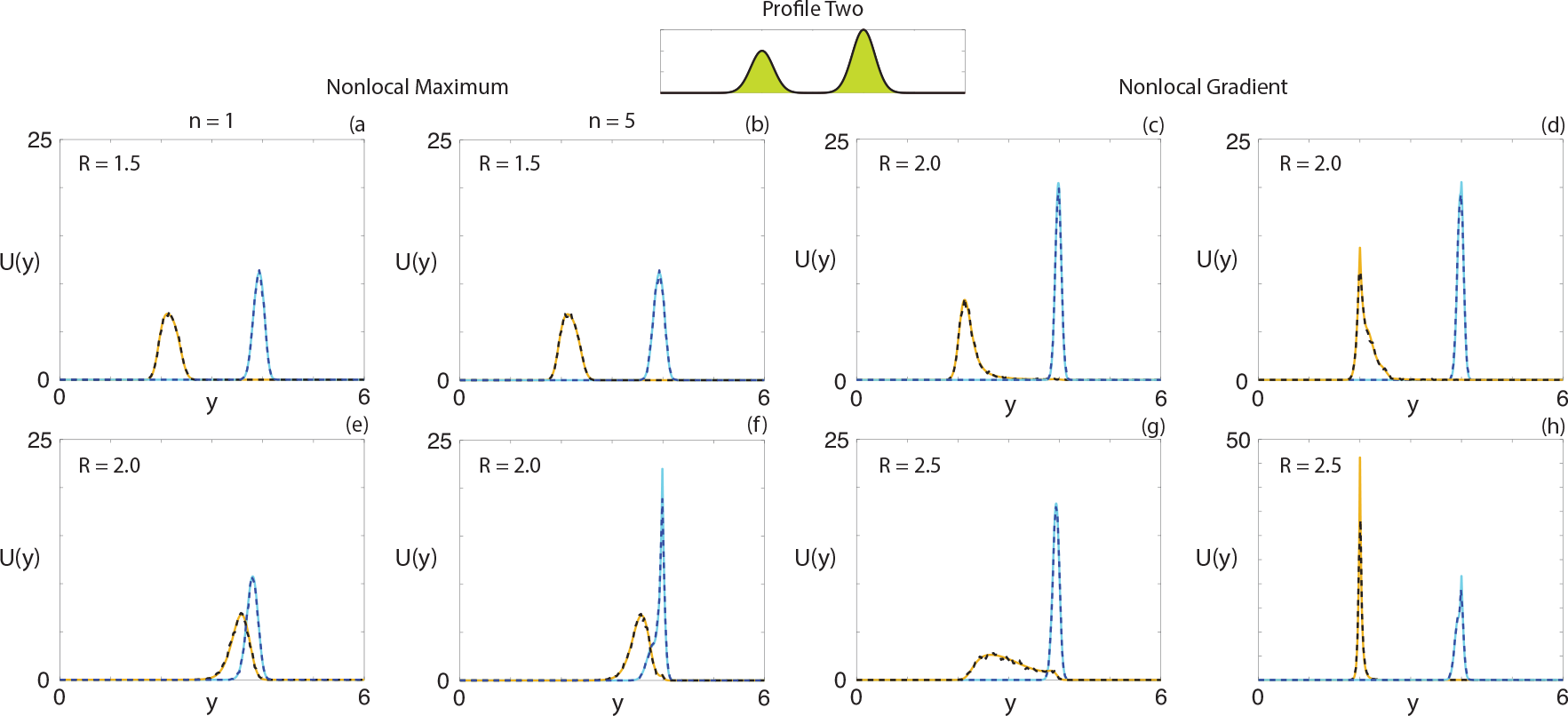
Comparison of different sensing radii and nonlinear order for the full DAD model (solid) and the average velocity-jump process (dashed), respectively, using (a,b,e,f) the nonlocal maximum model and (c,d,g,h) the nonlocal gradient model. Individuals are initially distributed on 1.8 ≤ *y* ≤ 2.2 (orange/black) and 3.8 ≤ *y* ≤ 4.2 (cyan/blue). The environmental cue corresponds to Profile Two. Parameters used are *t*_end_ = 1000, (a,b,e,f) *κ*_3_ = 3, (c,d,g,h) *κ*_2_ = 0.3, *M* = 10^5^, Δ*y* = 0.003, *τ* = 1, *s* = 0.01.

#### S2.2. Bodega Bay

Figure S.6 shows direction fields for Model (M2) with *n* = 1 and (M3) with *n* = 1 and *n* = 5.

### S3. Numerical methods

#### S3.1. Approximations and idealised study

Numerical solutions for the quasi-one-dimensional idealised studies of model (1) and its approximations are obtained using an operator splitting method. For the diffusion operator component of (1), the equation is spatially discretised using central finite differences, and temporally discretised in an implicit manner. The resulting system of tridiagonal algebraic equations is solved using the Thomas algorithm. For the advection operator component of (1), the equation is solved using the Lax-Wendroff method.

#### S3.2. Bodega marine reserve numerics

Numerical methods for solving the VJRW model and the DAD equation (1) for hilltopping on a natural terrain (Bodega Marine Reserve) have been adapted from our previous studies, e.g. [34], and we refer there for full details. Here we restrict to highlighting some specific points arising from the current paper.

Our terrain restricts to a rectangular study region of area 2.2 × 2.2 km^2^, encompassing a portion of the Bodega Marine Reserve, North California. As described in the main text, high spatial resolution LIDAR data was obtained from the United State Geological Survey’s “The National Map” project over the co-ordinate range for the study site. Elevation data is extracted at each data point and interpolation onto a regular grid is performed. This elevation data subsequently feeds into models (M1-M3) to generate the navigation fields used for orienteering decisions. Note that, for the VJRW, further interpolation of the elevation data is required to filter from the regular grid on which elevation data is required to the individual’s positon in 2D space. Numerical evaluations of the nonlocal models (M1-M3) also demand integration over a circular region of space, which is performed through converting to a polar coordinate system, discretising over the polar angle and radial coordinate and computing the integral via direct summation, again using interpolation to convert elevation data from the regular grid to that used to compute the integral. We remark that numerical explorations have been performed for a range of spatial discretisations with negligible change to solution behaviour.

**Figure S5:**
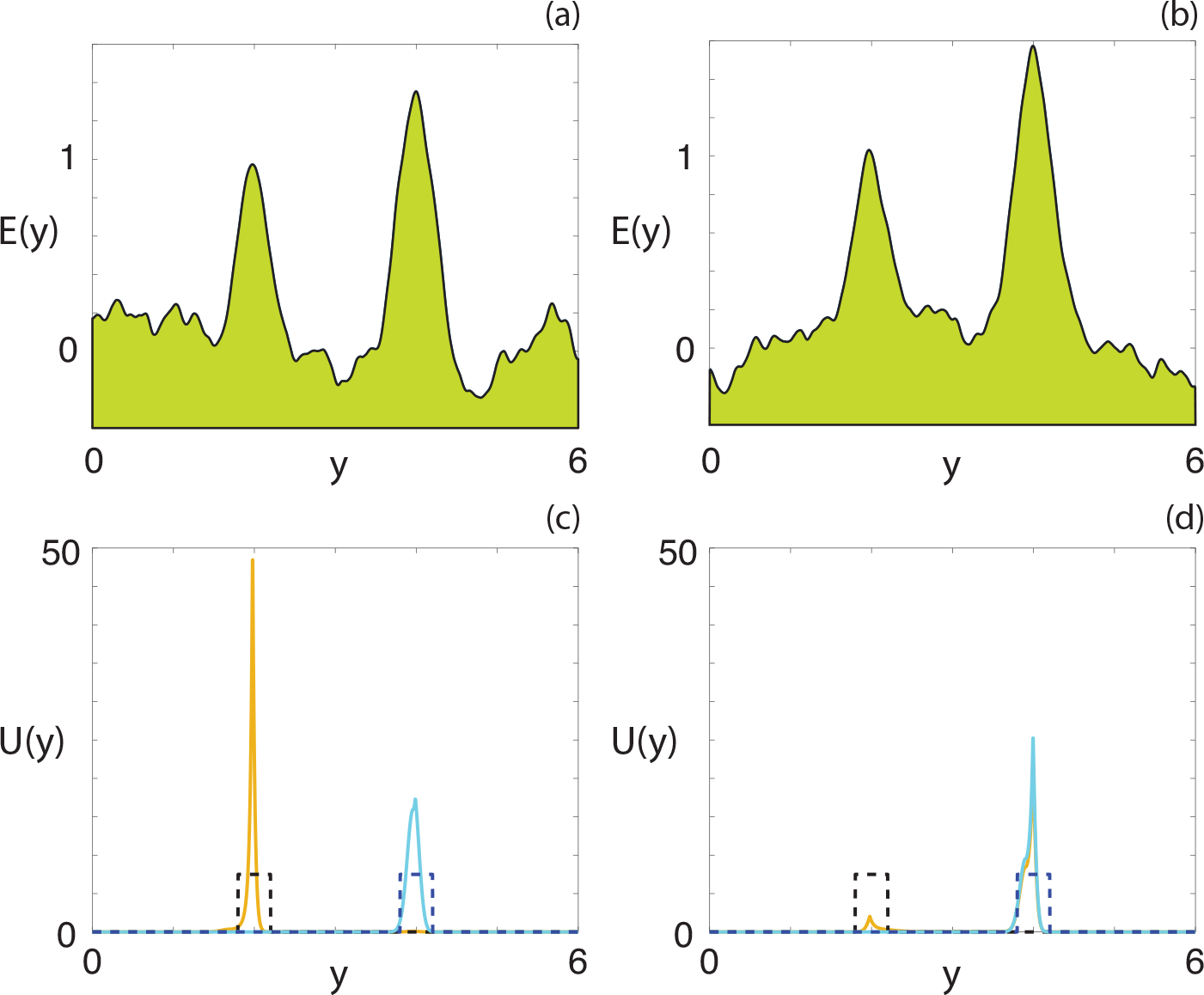
Comparison of two identically-prepared noisy environments, and the corresponding migration between peaks for the nonlocal maximum model. The initial condition for the orange (cyan) profile is given by the black (blue) dashed profile. The deterministic component environmental cue corresponds to Profile Two. Parameters used are *t*_end_ = 10^4^, *κ*_3_ = 3, *R* = 1.7, *n* = 5, Δ*y* = 0.003, *τ* = 1, *s* = 0.01.

**Figure S6:**
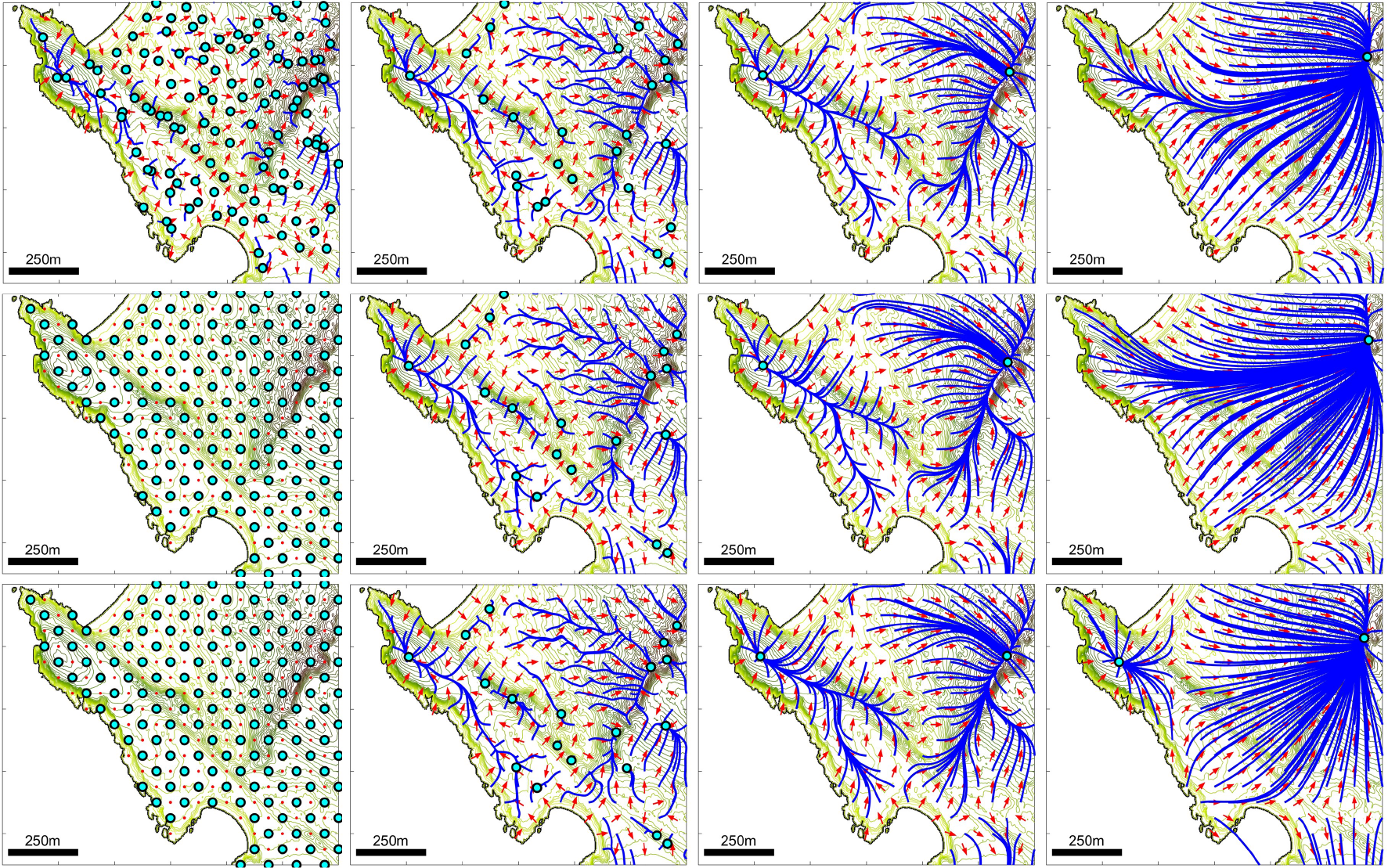
Representation of orientation information provided by **w**, where: (Top row) **w**is given by (M2) with *R* = 0 (M1), *R* = 25, *R* = 100 and *R* = 500 and *n* = 1; (Middle row) **w**is given by (M3) with *R* = 0, *R* = 25, *R* = 100 and *R* = 500 and *n* = 1; (Bottom row) **w**is given by (M3) with *R* = 0, *R* = 25, *R* = 100 and *R* = 500 and *n* = 1.

Special attention must be made concerning the movement of individuals at boundaries. In order to ensure that the full initial population remains in the study zone across the course of experiments, we implement a no-loss (zero-flux) boundary condition. Of course, this is a somewhat artificial construct, as the natural landmass extends beyond the artificial boundaries imposed by our study region and insects would certainly move to and fro according to elevation. However, (i) the no-loss choice facilitates our study by maintaining the same sized population within the study zone, and more realistic choices do not generate qualitatively different behaviour to that generated our model.

#### S3.3. Velocity-jump random walk

In the VJRW, *M* individuals are placed on the domain at locations 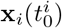, *i* = 1,…, *M* according to the appropriate initial distribution. To update the position of an individual at 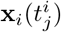, we sample a velocity 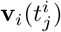 from the von Mises distribution with input parameters based on the navigation vector field at 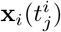. We next sample a runtime 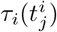 from a Poisson distribution with mean runtime *τ*. Combining these two sampled values, we update the location of the individual according to 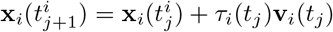 and update the time associated with that individual, 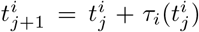. We repeat this process for all *M* individuals until each individual satisfies 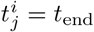 [34].

## References

[1] J. Alcock. Leks and hilltopping in insects. J. Nat. Hist., 21:319–328, 1987.

[2] A. Barnett and P. Moorcroft. Analytic steady-state space use patterns and rapid computations in mechanistic home range analysis. J. Math. Biol., 57:139–159, 2008.

[3] R. N. Binny, P. Haridas, A. James, R. Law, M.J. Simpson, and M.J. Plank. Spatial structure arising from neighbour-dependent bias in collective cell movement. PeerJ, 4:e1689, 2016.

[4] A. Buttenschoen, T. Hillen, A. Gerisch, and K. J. Painter. A space-jump derivation for non-local models of cell-cell adhesion and non-local chemotaxis. J. Math. Biol., 76:429–456, 2018.

[5] E. A. Codling, M. J. Plank, and S. Benhamou. Random walk models in biology. J. Roy. Soc. Interface, 5:813–834, 2008.

[6] B. J. Duistermars, D. M. Chow, and M. A. Frye. Flies require bilateral sensory input to track odor gradients in flight. Curr. Biol., 19:1301–1307, 2009.

[7] R. Eftimie. Hyperbolic and kinetic models for self-organized biological aggregations and movement: a brief review. J. Math. Biol., 65:35–75, 2012.

[8] P.R. Ehrlich and D. Wheye. “Nonadaptive” hilltopping behavior in male checkerspot butter-flies (euphydryas editha). Amer. Nat., 127:477–483, 1986.

[9] G. Estrada-Rodriguez, H. Gimperlein, and K. J. Painter. Fractional Patlak-Keller-Segel equations for chemotactic superdiffusion. SIAM J. Appl. Math., 78:1155–1173, 2018.

[10] W. F. Fagan, E. Gurarie, S. Bewick, A. Howard, R. S. Cantrell, and C. Cosner. Perceptual ranges, information gathering, and foraging success in dynamic landscapes. The American Naturalist, 189:474–489, 2017.

[11] E. A. Flaherty, W. P. Smith, S. Pyare, and M. Ben-David. Experimental trials of the northern flying squirrel (Glaucomys sabrinus) traversing managed rainforest landscapes: perceptual range and fine-scale movements. Can. J. Zool., 86:1050–1058, 2008.

[12] R. J. Fletcher, C. W. Maxwell, J. E. Andrews, and W. L. Helmey-Hartman. Signal detection theory clarifies the concept of perceptual range and its relevance to landscape connectivity. Landscape ecology, 28:57–67, 2013.

[13] R. Friedrich, F. Jenko, A. Baule, and S. Eule. Exact solution of a generalized Kramers-Fokker-Planck equation retaining retardation effects. Phys. Rev. E, 74:041103, 2006.

[14] J. L. Gould and C. G. Gould. Nature’s compass: the mystery of animal navigation. Princeton University Press, 2012.

[15] P. Grof-Tisza, Z. Steel, E. M. Cole, M. Holyoak, and R. Karban. Testing predictions of movement behaviour in a hilltopping moth. Anim. Behav., 133:161–168, 2017.

[16] T. Hillen. Transport equations with resting phases. European Journal of Applied Mathematics, 14(5):613–636, 2003.

[17] T. Hillen, K. Painter, and C. Schmeiser. Global existence for chemotaxis with finite sampling radius. Disc. & Cont. Dyn. Sys. B, 7:125–144, 2007.

[18] T. Hillen and K. J. Painter Dispersal, individual movement and spatial ecology: a mathematical perspective, chapter Transport and anisotropic diffusion models for movement in oriented habitats, pages 177–222. Springer, 2013.

[19] T. Hillen, K. J. Painter, A. C. Swan, and A. D. Murtha. Moments of von Mises and Fisher distributions and applications. Math. Biosci. & Eng., 14(3):673–694, 2017.

[20] C. D. Hopkins. Passive electrolocation and the sensory guidance of oriented behavior. In Electroreception, pages 264–289. Springer, 2005.

[21] N. L. Jeon, H. Baskaran, S. Dertinger, G. M. Whitesides, L. Van De Water, and M. Toner. Neutrophil chemotaxis in linear and complex gradients of interleukin-8 formed in a microfab-ricated device. Nature Biotech., 20:826–830, 2002.

[22] E.F. Keller and L.A. Segel. Initiation of slime mold aggregation viewed as an instability. J. Theor. Biol., 26:399–415, 1970.

[23] T. B. Kornberg and S. Roy. Cytonemes as specialized signaling filopodia. Development, 141:729–736, 2014.

[24] S. L. Lima and P. A. Zollner. Towards a behavioral ecology of ecological landscapes. Trends Ecol. & Evol., 11:131–135, 1996.

[25] K. V. Mardia and P. E. Jupp. Directional Statistics. Wiley, New York, 2000.

[26] K. McComb, D. Reby, L. Baker, C. Moss, and S. Sayialel. Long-distance communication of acoustic cues to social identity in African elephants. Anim. Behav., 65:317–329, 2003.

[27] A. M. Middleton, C. Fleck, and R. Grima. A continuum approximation to an off-lattice individual-cell based model of cell migration and adhesion. J. Theor. Biol., 359:220–232, 2014.

[28] A. Mogilner and L. Edelstein-Keshet. A non-local model for a swarm. J. Math. Biol., 38:534–570, 1999.

[29] P. R. Moorcroft and A. Barnett. Mechanistic home range models and resource selection analysis: a reconciliation and unification. Ecology, 89:1112–1119, 2008.

[30] J. D. Olden, R. L. Schooley, J. B. Monroe, and N. L. Poff. Context-dependent perceptual ranges and their relevance to animal movements in landscapes. J. Anim. Ecol., 73:1190–1194, 2004.

[31] F. W. J. Olver. NIST handbook of mathematical functions. Cambridge University Press, 2010.

[32] H. G. Othmer, S. R. Dunbar, and W. Alt. Models of dispersal in biological systems. J. Math. Biol., 26:263–298, 1988.

[33] H. G. Othmer and T. Hillen. The diffusion limit of transport equations II: Chemotaxis equations. SIAM J. Appl. Math., 62:1222–1250, 2002.

[34] K. J. Painter. Multiscale models for movement in oriented environments and their application to hilltopping in butterflies. Theor. Ecol., 7:53–75, 2014.

[35] K. J. Painter. Mathematical models for chemotaxis and their applications in self-organisation phenomena. J. Theor. Biol., Article in Press, 2018.

[36] K. J. Painter, J. M. Bloomfield, J. A. Sherratt, and A. Gerisch. A nonlocal model for contact attraction and repulsion in heterogeneous cell populations. Bull. Math. Biol., 77:1132–1165, 2015.

[37] C. S. Patlak. Random walk with persistence and external bias. Bull. Math. Biol., 15:311–338, 1953.

[38] R. Payne and D. Webb. Orientation by means of long range acoustic signaling in baleen whales. Ann. New York Acad. Sci., 188:110–141, 1971.

[39] G. Pe’er, S.K. Heinz, and K. Frank. Connectivity in heterogeneous landscapes: analyzing the effect of topography. Land. Ecol., 21:47–61, 2006.

[40] G. Pe’er and S. Kramer-Schadt. Incorporating the perceptual range of animals into connectivity models. Ecol. Model., 213:73–85, 2008.

[41] G. Pe’er, D. Saltz, and K. Frank. Virtual corridors for conservation management. Conservation biology, 19:1997–2003, 2005.

[42] G. Pe’er, D. Saltz, T. Münkemüller, Y. G. Matsinos, and HH. Thulke. Simple rules for complex landscapes: the case of hilltopping movements and topography. Oikos, 122:1483–1495, 2013.

[43] G. Pe’er, D. Saltz, H. H. Thulke, and U. Motro. Response to topography in a hilltopping butterfly and implications for modelling nonrandom dispersal. Anim. Behav., 68:825–839, 2004.

[44] K. Perlin. An image synthesizer. ACM Siggraph Computer Graphics, 19:287–296, 1985.

[45] B. Perthame. Transport equations in biology. Springer Science & Business Media, 2006.

[46] J. A. Prevedello, G. Forero-Medina, and M. V. Vieira. Does land use affect perceptual range? evidence from two marsupials of the Atlantic Forest. J. Zool., 284:53–59, 2011.

[47] J.A. Scott. Hilltopping as a mating mechanism to aid the survival of low density species. J. Res. Lepid, 7:191–204, 1968.

[48] O. Shields. Hilltopping. J. Res. Lepid, 6:69–178, 1967.

[49] J.P. Taylor-King, E.E. van Loon, G. Rosser, and S.J. Chapman. From birds to bacteria: generalised velocity jump processes with resting states. Bull. Math. Biol., 77:1213–1236, 2015.

[50] J. M. Teddy and P. M. Kulesa. In vivo evidence for short-and long-range cell communication in cranial neural crest cells. Development, 131:6141–6151, 2004.

[51] L. Todd, R. Poulin, R. Brigham, E. Bayne, and T. Wellicome. Pre-migratory movements by juvenile burrowing owls in a patchy landscape. Avian Cons. & Ecol., 2, 2007.

[52] YS. Wang and J.R. Potts. Partial differential equation techniques for analysing animal movement: A comparison of different methods. J. Theor. Biol., 416:52–67, 2017.

[53] L. S. Weilgart. The impacts of anthropogenic ocean noise on cetaceans and implications for management. Can. J. Zool., 85:1091–1116, 2007.

[54] P. A. Zollner and S. L. Lima. Landscape-level perceptual abilities in white-footed mice: perceptual range and the detection of forested habitat. Oikos, 80:51–60, 1997.

